# The evolutionary dynamics of organellar pan-genomes in *Arabidopsis thaliana*

**DOI:** 10.1101/2025.01.20.633836

**Authors:** Yi Zou, Weidong Zhu, Yingke Hou, Daniel B. Sloan, Zhiqiang Wu

**Author notes:** E-mails of other authors: Yi Zou, Weidong Zhu, Yingke Hou, Daniel B. Sloan.

## Abstract

In plants, comparative analyses of organellar genomes often depend on draft assemblies, with large-scale investigations into the complex structural rearrangements of mitochondrial genomes remaining scarce. Here, we conducted a comprehensive analysis of the dominant conformations and dynamic heteroplasmic variants of organellar genomes in the model plant *Arabidopsis thaliana*, utilizing high-quality long-read assemblies validated at read-level resolution from 149 samples. We found that mitochondrial and plastid genomes share common types of structural and small-scale variants driven by similar DNA sequence features. However, rearrangements mediated by repetitive sequences in mitochondrial genomes evolved so rapidly that they often became decoupled from other types of variants. Rare complex events involving elongation and fusion were also observed, contributing to the unalignable regions commonly found at the interspecies level. Additionally, we demonstrated that disrupting and rescuing organellar DNA maintenance could drive the rapid evolution of dominant mitochondrial genome conformations. Our study provides an unprecedentedly detailed view of the dynamics of organellar genomes at pan-genome scale in *A. thaliana*, paving the way to unlock the full potential of organellar genetic resources.

**Significance:** Plant mitochondrial genomes exhibit rapid rearrangements within species. Large-scale analyses using high-quality assemblies offer detailed insights into their evolutionary dynamics, paving the way for organellar genome engineering and functional research in the future.

## Introduction

Genetic variation forms the foundation of natural selection and adaptive evolution. In plants, genetic material is distributed across three compartments, the nucleus and the semi-autonomous organelles, mitochondria (mt) and plastids (pt). These two organellar genomes differ greatly in mutation rate and sequence features such as the size, repeat content, gene content, etc. (Wang et al., 2024b). Mutations in mt and pt genomes can directly impact plant growth and developmental processes. For example, pt genome variants have been associated with yellowing/whitening of leaves and different variegation patterns (Massouh et al., 2016; Malinova et al., 2021). Variegated leaves can help alleviate high light stress in the environment, while also holding significant horticultural ornamental value (Xu et al., 2011; Park et al., 2023). Variants in mt genomes have been shown to cause male sterility, curled or wrinkled leaves, and delayed seed germination (Chen and Liu, 2014; Virdi et al., 2016; Golin et al., 2020; Forner et al., 2023). Multiple copies of organellar genomes can exist within an individual or even a single cell, often with different structures or variants, which is known as heteroplasmy. In humans, nuclear genes with roles in DNA replication, recombination and repair (RRR) processes have been associated with mt heteroplasmy levels (Gupta et al., 2023). In plants, *MutS homolog 1* (*MSH1*) is one of the most well-studied genes involved in organellar RRR processes (Martínez-Zapater et al., 1992; Sakamoto et al., 1996; Abdelnoor et al., 2003; Shedge et al., 2007; Arrieta-Montiel et al., 2009; Davila et al., 2011; Zou et al., 2022; Sloan et al., 2024). The *A. thaliana MSH1* gene accelerates “heteroplasmic sorting” across generations potentially through gene conversion between organellar genome copies, with the sorting of pt genome variants occurring rapidly even in the absence of *MSH1* (Broz et al., 2022, 2024). However, the contribution of individual-level heteroplasmy in shaping the distinct evolutionary dynamics of organellar genomes remains unclear, especially concerning the less-explored structural variants (SVs) of mt genomes.

Despite their significance, studies of plant mt genomes lag behind those of pt and nuclear genomes (Wang et al., 2024b). The challenge of sequencing and assembling plant mt genomes is primarily due to their abundant repeats (large, ≥ 1 kb; intermediate-sized, 50 bp–1 kb; small, < 50 bp), which play active roles in RRR processes (Christensen, 2013; Gualberto and Newton, 2017). These repeats often cause mapping issues and are not fully addressed by existing tools for organellar genome analysis, such as PMAT (Bi et al. 2024) and TIPPo (Xian et al., 2025b). Moreover, tools for evaluating the completeness, structural conformation, and base-level accuracy of plant mt genomes are not well-established. Under certain stress conditions or in mutants lacking key organellar RRR machinery, there can be elevated variant frequencies, further complicating the assembly of organellar genomes and variant quantification (Gualberto and Newton, 2017; Mackenzie and Kundariya, 2019). In organellar genome editing studies, researchers may overlook structural heteroplasmies that might explain phenotypic differences among lines by relying on short-read sequencing or PCR around the target regions, which fails to capture larger SVs (Arimura et al., 2020; Takatsuka et al., 2022; Forner et al., 2023; Zhou et al., 2024). Another challenge in studying organellar genomes is the existence of large mt and pt-derived sequences in the nucleus (NUMTs and NUPTs, respectively), which can exhibit high sequence similarity to organelle genomes (Ma et al., 2020). Because there are fewer copies of mtDNA than ptDNA per cell, mt genome analysis is more susceptible to artefactual effects of NUMTs. When quantifying variant frequencies in organellar genomes, NUMTs and NUPTs can often be distinguished by the differences in their sequences. However, for some long identical regions, distinguishing true organelle genome heteroplasmies is challenging (Fields et al., 2022). Although differential DNA methylation levels present a potential solution, achieving this at read-level resolution remains difficult (Zhong et al., 2024). To address these challenges with short-read technologies, we previously developed a pipeline based on PacBio high-fidelity (HiFi) long-read sequencing that can quantify global patterns of SVs in mt genomes (Zou et al., 2022). We also generated draft mt genome assemblies representing the pseudo-master circles of seven *A. thaliana* accessions using published PacBio continuous long read (CLR) data, which have higher error rates than HiFi reads. However, the correctness of these assemblies and the interference from NUMTs/NUPTs were not examined. Moreover, we did not explore the intraspecific evolution of organellar genomes based on high-quality assemblies at large scales in *A. thaliana*.

In this study, we aim to quantitatively examine the intraspecific organellar genome variation in *A. thaliana*, with a particular focus on the structural dynamics of mt genomes. We collected published HiFi datasets consisting of 149 samples from 140 distinct accessions. We assembled and validated the accuracy of mt and pt genome assemblies using a custom pipeline that extends and refines the functionalities of hifisr (Zou et al., 2022). By providing sample-specific snapshots at read-level resolution, the new pipeline was able to obtain a master conformation representing the most frequent structural and small-scale alleles for each sample, while still recovering all low-frequency variants including partial alignments derived from NUMTs and NUPTs. For each accession, the organellar genomes exhibited clear dominant conformations, except in regions of large repeats and long homopolymers. By comparing the dominant conformations across *A. thaliana* accessions, we found that the mt and pt genomes exhibited similar types of structural rearrangements driven by tandem and non-tandem repeats. However, mt genomes possessed an additional class of intermediate-sized repeats that underwent rearrangements at a higher frequency. In rare instances, we observed repeat elongation and fusion in certain accessions, a phenomenon that is increasingly common at the interspecies level. We further demonstrated that disrupting and rescuing a single nuclear-encoded gene, *MSH1*, enabled the generation of different structural versions of mt genomes, while the structure of pt genomes remained resistant to such changes. These findings provide insights into the connections between organellar genome variation at individual and population levels in *A. thaliana*, paving the way to unlocking the full potential of organellar genetic resources.

## Results

### Validating organellar genome assemblies by variant frequency estimation

We collected published PacBio HiFi datasets of 149 samples of *A. thaliana* (five for Col-0, three for Etna-2, two each for Ms-0, St-0, Tanz-1, and 135 for other accessions), and developed a three-stage pipeline for assembling the organelle genomes and validating their accuracy (Figure 1a). In the Assembly stage, we first generated metaFlye assembly graphs using bait-enriched reads for each sample. We found that nearly all the assembly graphs had zero to two unresolved large repeats and could be further resolved to get the linear representations of mt and pt genomes except for a few samples with high proportions of NUMT reads (Figure S1). In the next stage, we polished rotated versions of the linear genomes using a custom Python script (see Methods). The rotation step ensured the successful correction of assembly errors at the boundaries of large repeats and was also critical for downstream variant calling (Figure S2). In the final Validation stage, we first collected all the reads that fully aligned to the mt and pt genomes, which revealed significant variation in total reads and bases across samples (Figure 1b, c; Table S1). Then, we checked the completeness of read coverage, estimated the frequencies of all the identified variants, and corrected errors in cases where the reported reference allele did not have the highest frequency (Figure 2a, b). The final linear genomes were aligned to Col-0 references based on BLASTn results, with mt genome sizes ranging from 337,346 to 368,977 and pt genomes sized ranged from 154,172 to 154,689 (Table S1). The mt genomes were further categorized based on copy numbers of large repeat pairs, following the naming scheme of previous studies (See details in Methods). Class 1 genomes contain two copies of Large1 and two copies of Large2, Class 2 genomes contain one copy of Large1 and two copies of Large2, and Class 3 genomes are a subgroup of Class 2 with two copies of Large3 (a pair of 1038-bp repeats present in only seven accessions; Figure 1d; Table S1).

**Figure 1.**
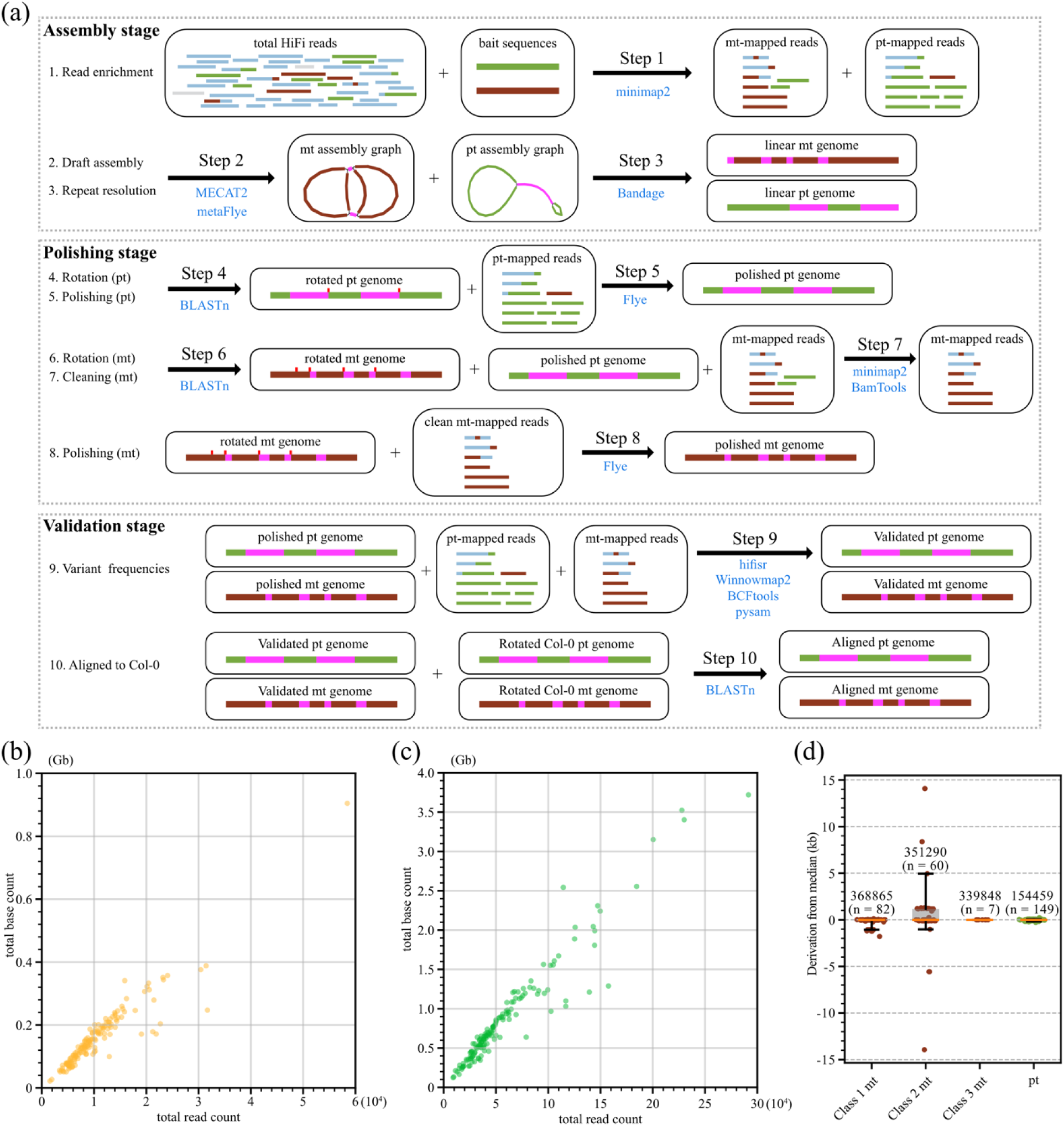
An overview of the bioinformatic pipeline based on HiFi reads. (a) A three-stage pipeline for assembling the organelle genomes and validating their accuracy. (b, c) The 149 samples contained various amount of reads that fully aligned to the (b) mt and (c) pt genomes. (d) The numbers and lengths of pt genomes and three classes of mt genomes.

**Figure 2.**
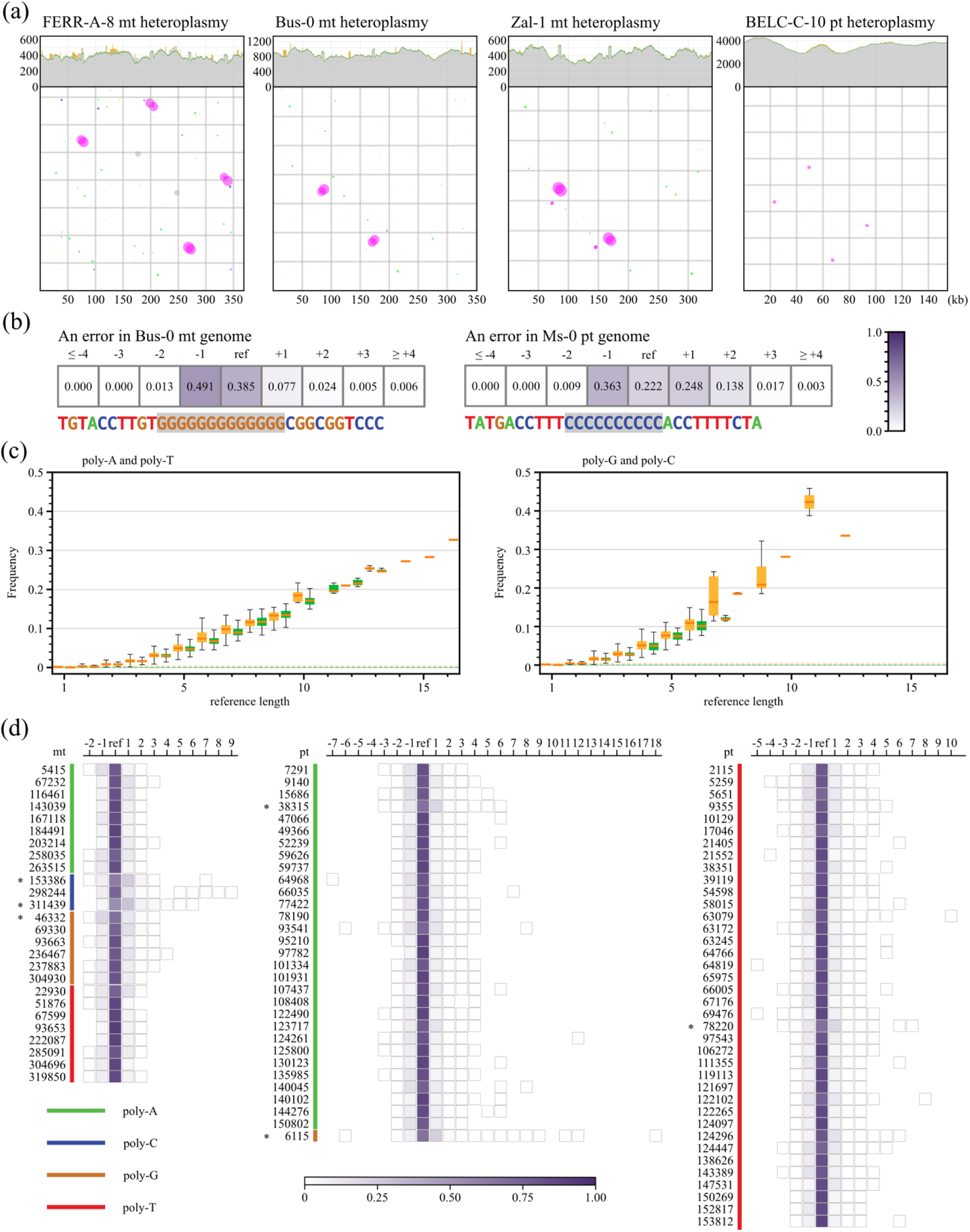
Dominant structural and small-scale variants support the accuracy of organellar genome assemblies. (a) Coverages (top) of HiFi reads in representative samples of pt genomes and three classes of mt genomes. Grey areas indicate fully aligned reads, orange areas indicate partially aligned reads, and the green line indicate reads with no rearrangements plus one-rearrangement reads mediated by large repeats (Large1, Large2, and IRab). Frequencies (bottom) of one-rearrangement reads in each sample. Colors represent categories of alignment overlap length (AO length) as in Zou et al. (2022). (b) Examples of errors where the frequencies of the reference allele were not the highest. (c) Variant frequencies at homopolymers increased with the reference length in mt (orange) and pt (green) genomes of Col-CEN. (d) Allele frequencies of different lengths at long homopolymers (reference length ≥ 10 bp) in Col-CEN.

According to our analysis, the frequencies of SVs were generally low except at the Large1 and Large2 repeat pairs in mt genomes, and the large inverted repeats (IRab) in pt genomes (Figure 2a and Figure S3a). Relative to single-copy sequences, there was only a slight increase in SV frequencies at the 1038-bp Large3 repeats, some intermediate-sized (50-1000 bp) non-tandem repeats, and some tandem repeats with a long unit length or a high copy number (Figure S3b-d). The estimated variant frequencies at some homopolymers (i.e. tracts of poly-A, poly-C, poly-G, and poly-T) were even higher than intermediate-sized repeats and occurred repeatedly in multiple samples. This could be due to the intrinsic instability of homopolymers, which increases rapidly with the reference length (Figure 2c). For long homopolymers (reference length ≥ 10 bp), we chose the most frequent allele as the reference length. We found that the most frequent length variants were 1-bp InDels, and the length distribution was skewed toward insertions (Figure 2d). Some of these long homopolymers were so unstable that different samples of the same accession (e.g., Col-0, Etna-2, and Ms-0) had a different dominant length (Figure S4; Table S2). However, determining variant frequencies at homopolymers is challenging because we cannot distinguish true biological variants from sequencing errors in HiFi reads. Another challenge was that high noise derived from NUMTs and NUPTs could introduce errors during genome assembly and polishing in some accessions. For example, the middle and ends of the Lu-1 mt genome exhibited an artefactual signal of two high-frequency SVs and increased read depth corresponding to four-copy regions in a 774,161-bp NUMT in the nuclear genome (Figure S5; Table S3). In this case, we skipped the polishing step and selected the proportion of zero-rearrangement reads to validate the correctness of the mt genome assembly. In sum, we have generated high-quality organellar genomes representing the dominant structural forms and the most frequent alleles of small-scale variants in 140 *A. thaliana* accessions.

### Exploring the structural variability of organellar genomes across different accessions

We analyzed the structural variability of organellar genomes by combining the results of both pairwise and multi-accession comparisons. Here, we used a BLASTn-based Python script in hifisr to identify structural rearrangements between two assemblies (see Methods). Compared with Col-0, the pt genomes in each accession contained up to two large InDels, which were often associated with copy number variants (CNV) of tandem repeats, microhomology-mediated end-joining (MMEJ) events (defined here as being mediated by small direct repeats of < 50 bp in size), and/or non-homologous end-joining (NHEJ) events (defined here as involving breakpoints with no more than 1 bp of shared sequences) (Table S4). In contrast, there was a range of repeat-mediated structural rearrangements for mt genomes. The Large1, Large2, B, C, Q, H, I, and L repeats were the most frequently involved in these rearrangements.

To summarize relationships between repeat-mediated mt SVs, we “flipped” sequences between inverted repeats starting from the Col-0 conformation in an iterative process until no new conformations could be generated. Such flipping operations may transform direct repeats in one conformation into inverted repeats in the next, and vice versa. Using only the repeats Large1, Large2, B, C, and Q, we generated a fully connected network of 21 possible conformations (nodes) (Figure 3a). For simplicity, the nodes connected by only inverted repeats Large1 and/or Large2 could be further flipped to keep only one node for each conformation group. For simplicity, Class 1 mt genomes were further assigned to twelve different conformation groups (Figure 3a; with details in Table S5). Similarly, Class 2 (twelve different conformation groups) and Class 3 (one conformation group) mt genomes were connected to this network by additional repeats (Figure 3a; Table S5). In addition to repeat-mediated rearrangements, we analyzed other structural changes by comparing the accessions within each group (Table S5). For example, we found rearrangements mediated by other non-tandem repeats in a few accessions, including the inverted repeats Large3 (Sha, Altai-5, Kar-1, Kyr-1, Nov-02, Sus-1, Zal-1), HNR1 (in Belmonte-4-94 and Ru-2), A (in Elh-2), T (in Elh-2), and D (in Hum-4, MONTM-B-16, St-0, T850, and Pyl-1); the direct repeats LL (in L*er*-0 and TueSB30-3); and a pair of 21-bp small repeats with one mismatch (in Kz-9). Similar to pt genomes, CNVs of tandem repeats, along with MMEJ and NHEJ events also contributed to large InDels in mt genomes (Table S5). These results suggest that, although rearrangements mediated by intermediate-sized repeats and non-tandem repeats were less common within a single accession, they were frequently involved in organellar genome evolution across different accessions.

**Figure 3.**
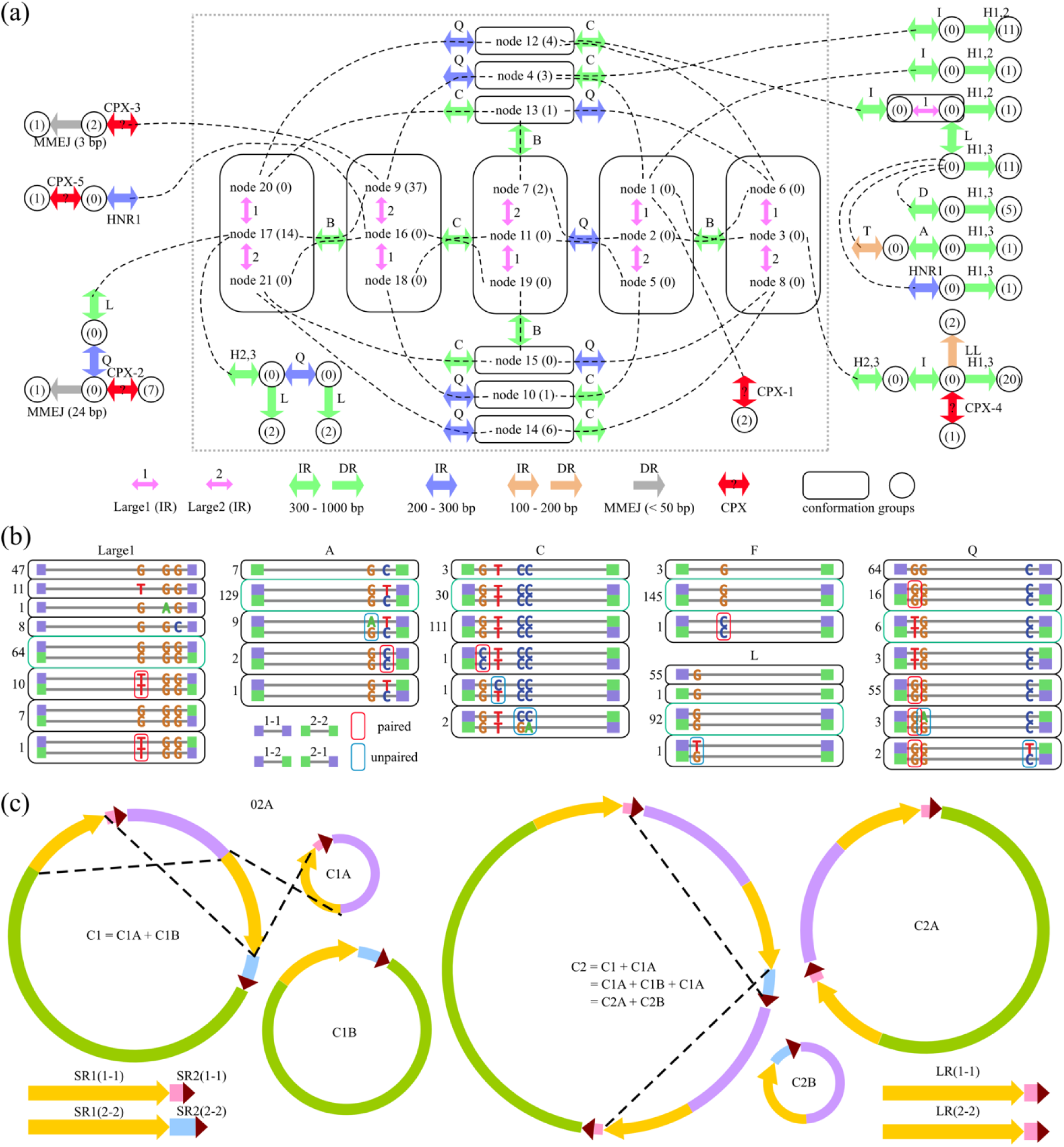
Repetitive sequences as primary drivers of structural variation in organellar genomes. (a) A network indicating the mt genome conformation groups of 140 *A. thaliana* accessions. Adjacent conformations groups were connected by direct repeats (single-head arrows), inverted repeats (double-head arrows with a repeat name), and complex (CPX) events (red double-head arrows with a question mark). Conformation groups for Class 1 mt genomes were in dashed rectangle. Numbers in parentheses indicate the count of accessions in each conformation group (b) Examples of mutation and gene conversion events in different repeats. The numbers on the left represent the count of accessions with the specified bases. The cases that were the same as Col-0 were in green boxes. (c) A possible model showing the intermediate conformations that lead to the fusion of shorter repeats.

Mutations reduce sequence similarity between repeat copies, whereas gene conversion events have the opposite effect. The lower frequency of rearrangements mediated by intermediate-sized repeats was further supported by the observation that large repeats always exhibited perfect sequence identity, whereas the two copies of intermediate-sized repeat groups, such as A, C, L, Q, were perfectly identical in some accessions but exhibited sequence differences in others (Figure 3b; Table S6 and S7). Thus, as expected, gene conversion events occur much more frequently for large repeats than for intermediate-sized repeats.

We also identified five complex (CPX) presence/absence changes involving sequences unalignable by BLASTn (Figure 2a and Figure S6; Table S8). Three of these could be explained by elongation of existing repeat groups. In CPX-1 (observed in Col-0 and Est-0), repeat HNR7-2 formed by a 60-bp repeat (Figure S7a). In CPX-4 (observed in Geg-14), a pair of 19,653-bp repeats formed by extending repeat C (Figure S6c). In CPX-5 (observed in Belmonte-4-94), Large2 was elongated to 4271 bp (Figure S7b). We also identified three cases where longer repeats formed by the fusion of closely located shorter repeats. Specifically, repeat W fused with repeat HNR27 in Belmonte-4-94 (Figure S7c), repeat H fused with repeat HNR36 in Cas-0 and Cat-0 (Figure S7d). In five accessions (Evs-0, Evs-12, IP-Ini-0, Met-6, Sah-0), the sequence between repeat MM-1 and repeat J-1 was replaced with that between repeat MM-2 and repeat J-2, leading to the fusion of repeats MM and J (Figure S7e). We hypothesize that these elongation or fusion events could happen through a series of intermediate conformations resulting from recombination events mediated by both shorter repeats (Figure 2c). Although the CPX-2 and CPX-3 events also involved accession-specific repeats, they were hard to explain due to the lack of obvious intermediate steps (Figure S6a, b). Across species, the mt genomes were highly rearranged and contained even more unalignable regions, making it impossible to track the intermediate steps, as observed for comparisons with outgroup species such *A. lyrata* and *Capsella bursa-pastoris* (Figure S8; Table S9). Taken together, our analyses offer a species-wide dynamic perspective on the structural variability of organellar genomes mediated by both tandem and non-tandem repeats in *A. thaliana*.

### Integrated pan-genomic analysis of the contrasting evolutionary dynamics of mt and pt genomes

To get an integrated pan-genomic understanding of mt and pt genomes in these *A. thaliana* accessions, we used BCFtools to identify genome-wide small-scale variants, including single-nucleotide variants (SNVs) and small InDels (see Methods). We conducted both pairwise and multi-accession comparisons, describing all the SVs and small-scale variants using the same Col-0 reference coordinates (Figure 4a; Table S10 and S11). Most of the small-scale variants in mt and pt genomes were limited to a small number of samples, and nearly half of them were private to a single accession (Figure S9; Table S10 and S11). Compared with the pt genomes, the mt genomes of *A. thaliana* had a higher ratio of non-coding regions, a higher GC content, a higher repeat content, and a more than four-fold lower density of small-scale variants (Figure 4a). The variants near SVs or within low-complexity AT-rich regions were very difficult to interpret. An example was the region around 141 kb in pt genomes, which exhibited very complex repeat patterns. However, we found that most of the closely located SNVs and/or small InDels in the same accession could be explained by the underlying DNA sequence features, including (1) gene conversion events between different copies of non-tandem repeats (Figure S10), (2) micro-inversion events mediated by partial palindromic repeats (Figure S11 and S12), and (3) some rare events involving the extension of repeat borders (Figure S13). These cases were more appropriately explained by a single mutational event that involved the changes of multiple bases. We also found that some mutations were better explained if we used a different reference accession for pairwise comparison, which suggested that information from multiple accessions could help to reveal the stepwise changes of some complex regions (Figure S14).

**Figure 4.**
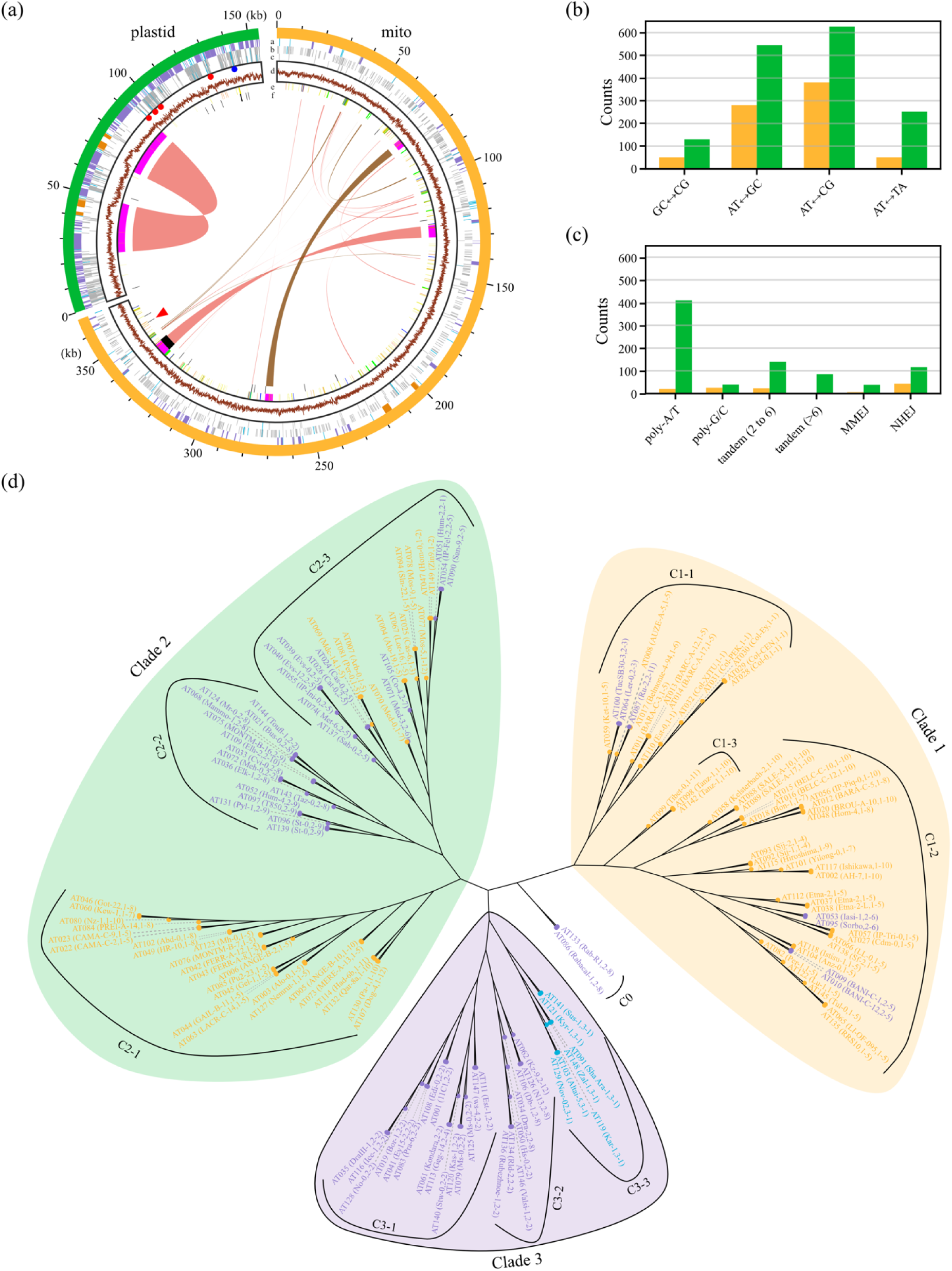
The structural and small-scale variants in organellar genomes of 140 *A. thaliana* accessions. (a) Circos plot shows the SVs, SNVs and small InDels in mt and pt genomes of all samples. Different tracks from a to f indicate gene annotations, SNVs, small InDels, GC content, repeat annotations, SVs that were not associated with non-tandem repeats. The chords indicate SVs that were associated with non-tandem repeats. The four red circles indicate AT-rich regions, and the blue circle indicates the region with complex repeat patterns in pt genomes. The red arrowhead indicates the 1105-bp unalignable region in mt genomes, which was absent in only the Col-0 and Est-0 accessions. No variants were found in this region. (b, c) Counts of different variant types of (b) SNVs and (c) small InDels. (d) An unrooted tree with three clades based on manually curated high-confidence SNVs in pt genomes. The colored circles at the tips indicate different classes of mt genomes (orange for Class 1, purple for Class 2, and blue for Class 3. The colored labels indicate sample indexes with accession names and mt genome conformation groups in parentheses.

We next analyzed the types and effects of small-scale variants based on DNA sequence features. Although the SNV frequencies were higher in pt genomes, we found that the most frequent types of SNVs were AT↔CG transversions in both genomes, followed by AT↔GC transitions (Figure 4b). The pt genomes also had higher frequencies of all types of InDels than mt genomes, especially the length variants mediated by poly-A and poly-T tracts (Figure 4c). Of the 873 variant sites (762 SNVs and 111 InDels) in mt genomes, only 27 SNVs (11 synonymous, 15 missense, 1 nonsense) were located in coding regions (Table S10 and Table S12). Of the 2129 variant sites (1525 SNVs and 605 InDels) in pt genomes, 459 SNVs (251 synonymous, 207 missense, 1 nonsense) and 12 InDels (11 non-frameshift, 1 frameshift) were located in coding regions (Table S11, S12, S13). Compared to the proportions of coding regions in mt (10.7%) and pt (58.6%) genomes, the lower proportions (3.1% for mt genomes and 22.1% for pt genomes) of variants in these regions suggests strong negative selection against deleterious mutations. These mt results are roughly consistent with a previous study based on the 1001G Illumina dataset (Wu et al., 2020a).

At the intraspecific level, SNVs and SVs mediated by small repeats occurred infrequently, whereas SVs mediated by intermediate-sized repeats and length variants of homopolymers occurred repeatedly in different accessions. In addition, the structure of the mt genomes evolved quickly compared with pt genomes. To further elucidate the contrasting evolutionary features of the mt and pt genomes, we began by constructing an unrooted tree using manually curated high-confidence pt SNVs. The results indicated that the 149 samples could be grouped into three main clades (Figure 4d), consistent with findings from a recent study based on short-read mapping of 1541 accessions (Theeuwen et al., 2024). After mapping the mt genome conformation groups (hereafter mt groups) to this tree, we found that most branches contained multiple mt groups (Figure 4d; Table S14). This inconsistency was observed even among closely related accession pairs in some cases (Figure 4d). In certain pt subclades, one or two mt groups dominate, potentially reflecting their origins. For example, the seven accessions of C3-3 located in eastern Central Asia, which belonged to the 3-1 mt group containing Large3 repeats (Figure 4d and Figure S15). The mt genomes of all samples had two copies of Large2, but Large1 repeats were included in only Class 1 mt groups. In most subclades, the samples either included the Large1 repeats (C1-3 and C2-1) or did not (C0, C2-2, C3-1, C3-2 and C3-3). However, for subclade C2-3, there were almost equal numbers for both cases (Figure 4d). Although all samples of this subclade are from the Iberian Peninsula (Figure S15 and Table S14), their mt groups showed no correlation with the reported nuclear genetic groups (Wlodzimierz et al., 2023). Therefore, caution is warranted when interpreting mt groups as phylogenetic signals, given the recurrent nature of SVs mediated by large and/or intermediate-sized repeats. Taken together, our analyses provide an integrated understanding of the intraspecific organellar genomic variants in *A. thaliana*, with an emphasis on the contrasting evolutionary dynamics of mt and pt genomes.

### Saltatory changes in mt genomes via disruption and rescue of the nuclear-encoded *MSH1* gene

Transient disruption of MSH1 has been identified as a plausible explanation for the rapid rearrangements observed in mt genome evolution, as supported by extensive studies (Martínez-Zapater et al., 1992; Sakamoto et al., 1996; Abdelnoor et al., 2003; Shedge et al., 2007; Arrieta-Montiel et al., 2009; Davila et al., 2011; Zou et al., 2022). To further explore this idea, we transformed *A. thaliana msh1* mutants with the *MSH1pro-MSH1gDNA-GFP* construct to generate transgenic plants and sequenced eight T5 individuals from four different rescue lines, designating *msh1*-RL (#5-1, #5-8, #6-12, #10-5). AlphaFold 3 (Abramson et al., 2024) predictions suggest that the GFP fusion at the C-terminus of MSH1 should not interfere with its DNA binding ability (Figure S16). Consistently, all eight individuals exhibited low SV frequencies across the mt genomes (except at large repeats) and a few nearly homoplasmic sites when compared to the original Col-0 mt genome (Figure S17a-g). Therefore, restoring *MSH1* functions appeared to suppress the high levels of heteroplasmic variation that is typical of *msh1* mutants (Arrieta-Montiel et al., 2009; Davila et al., 2011; Zou et al., 2022). Based on the simple metaFlye assembly graphs for these rescue lines, we successfully obtained their dominant mt genome conformations (Figure 5a). The accuracy of these assemblies was supported by the frequencies of SVs and small-scale variants using our pipeline (Figure S17i-p). Compared with Col-0, the ‘evolved’ mt genomes in the rescue lines had 2 to 5 SVs mediated by repeat Large1, Large2, MMJS, D, H, O, Y, and a tandem repeat (Figure 5a; Table S15). In total across the four lines, we identified one SNV, one MMEJ event mediated by a 13-bp micro-homology, a few 1-bp InDels at long homopolymers, one repeat extension event (Large1 plus 5 bp), and many gene conversion events (at repeat MMJS, A, G, H, N, O, P, V, W, Y, GG, and a 14-bp micro-homology), excluding those ambiguous variants near SVs (Table S16). In *msh1*-RL #6-12, the repeat A pair became perfectly identical in sequence due to gene conversion, and Large1 became 1217-bp shorter due to the 1610-bp deletion mediated by the recombination between Y-1 and Y-3. More interestingly, we found that the dominant conformation of mt genomes was not circular in *msh1*-RL #5-8 (Figure 5a, b). Derived from the same T1 parent, the circular mt genome in *msh1*-RL #5-1 and the non-circular mt genome in *msh1*-RL #5-8 shared two SVs (mediated by repeat H-1/H-3 and Large2), three 1-bp InDels at homopolymers, and all the gene conversion events at repeat O, V, GG Y, G (Table S15 and S16). In *msh1*-RL #5-1, the mt genomes had two additional SVs (mediated by repeat MMJS and Large1). Whereas in *msh1*-RL #5-8, the recombination events mediated by MMJS and Large1 were nonreciprocal, resulting in the non-circular mt genome with a 7720-bp deletion with only a 494-bp non-redundant sequence (Figure 5c). By examining the sequences at the links of different contigs, we found that one recombination breakpoint was located inside of MMJS (77-bp away from the border), while the other recombination breakpoint was located at the border of Large1 with a 5-bp extension (Figure 5b). In contrast to the mt genomes, the pt genomes of *msh1*-RL (#5-1, #5-8, #6-12, #10-5) were all 154,478-bp long and showed no structural variation (Figure S18). Besides, we identified two SNVs in #5-1 and #5-8, and five to seven 1-bp InDels at long homopolymers in the pt genomes for each line (Table S17). Taken together, these results demonstrate that, by disrupting and then rescuing the *MSH1* gene, different heteroplasmic versions of the mt and pt genomes can be generated and then fixed in a homoplasmic state after restoration of *MSH1* and that repeats and long homopolymers tend to be the main players in this kind of evolution.

**Figure 5.**
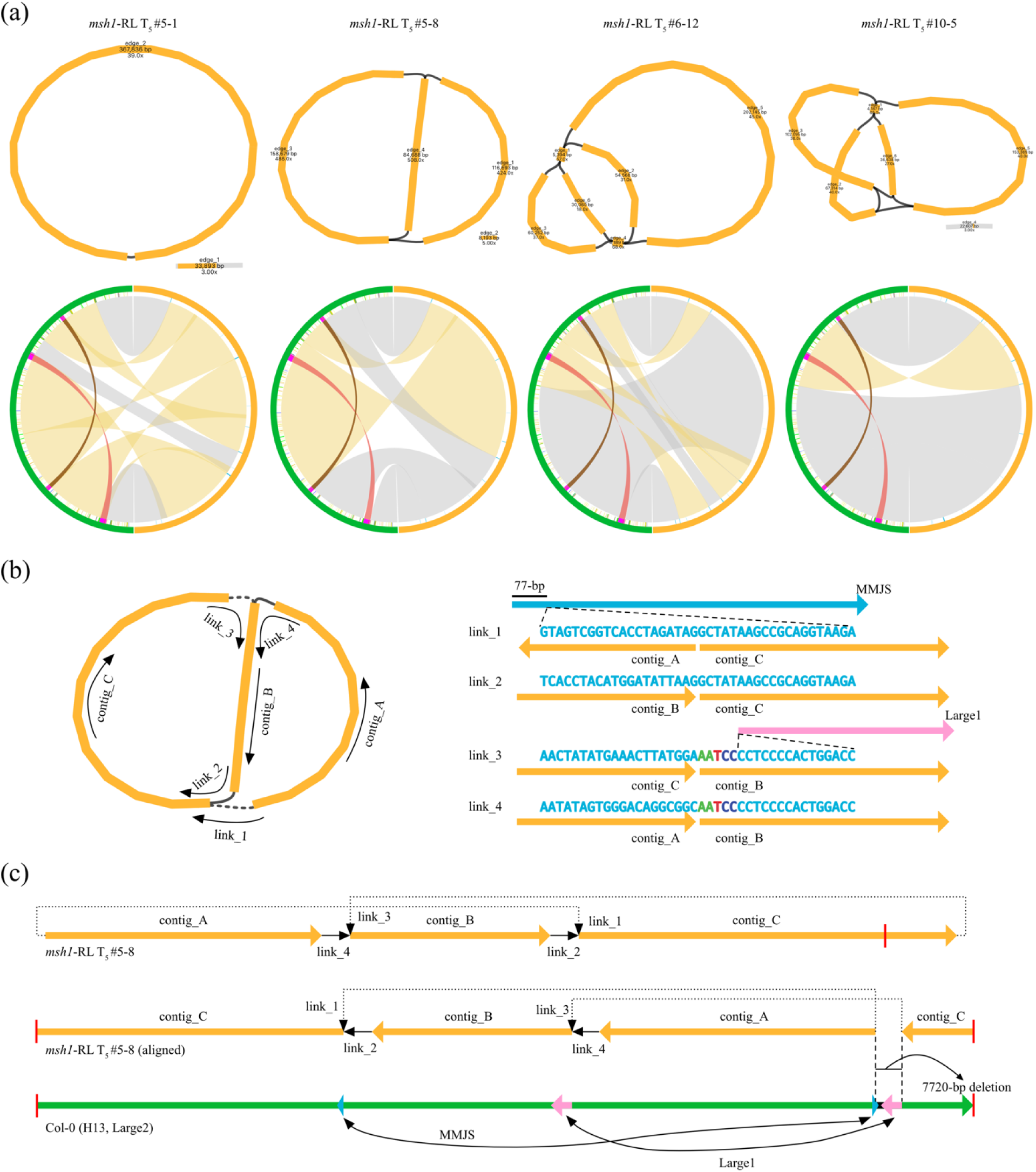
Structural comparison of dominant mt genome conformations in *MSH1* rescue lines. (a) Mitochondrial genome assembly graphs (top) of four rescue lines of *msh1* mutants. Circos plots (bottom) show the SVs and small-scale variants compared with the original Col-0 mt genome. (b) The dominant conformation (left) of mt genomes was not circular in *msh1*-RL #5-8. The sequences (right) at the links of adjacent contigs were shown. (c) The non-circular mt genome were generated by nonreciprocal recombination events at repeats MMJS and Large1. Top, linear representation of the assembly graph in *msh1*-RL #5-8. Middle, the reverse-complemented and rotated view of the same linear representation. Bottom, linear representation of Col-0 mt genome conformation after flipping repeats H1,3 and Large2.

We next investigated the organellar genome mutational landscape within individuals across different genetic backgrounds. Compared with the F3 individuals sequenced previously (Zou et al., 2022), the F2 individuals of *msh1* T-DNA insertional mutants (hereafter referred to as *msh1-*T-DNA) exhibited a higher ratio of reference conformations across the mt genome, including the most unstable region (F_2_: 43.58–49.90% reference; versus F_3_: 0.00–18.75%) around Large1-2 in Col-0 mt genome (Figure 6a and Figure S19). For small-scale variants, the frequency distribution in F2 *msh1* mutants was very similar to wild type, except for a slight rise in repeat-related SNVs in mt genomes (Figure 6b). In F3 *msh1* mutants, many sites of SNVs and small InDels rose in frequency; while in the rescue lines, the distribution returned to a low-heteroplasmy state with some variants becoming (nearly) fixed (Figure 6b). Despite using fully aligned reads, it was important to note that the variant frequencies still contained signals from the large NUMT on Chr2 (Figure S20). For pt genomes, the frequencies of small-scale variants in F2 *msh1* mutants were indistinguishable from wild type, while frequency rise and fixation were observed in F3 and rescue lines, respectively (Figure 6c). Consistent with the slower rate of structural evolution in the pt genome, the frequencies of pt SVs within *msh1* individuals were only slightly increased at sporadic locations (Figure S21).

**Figure 6.**
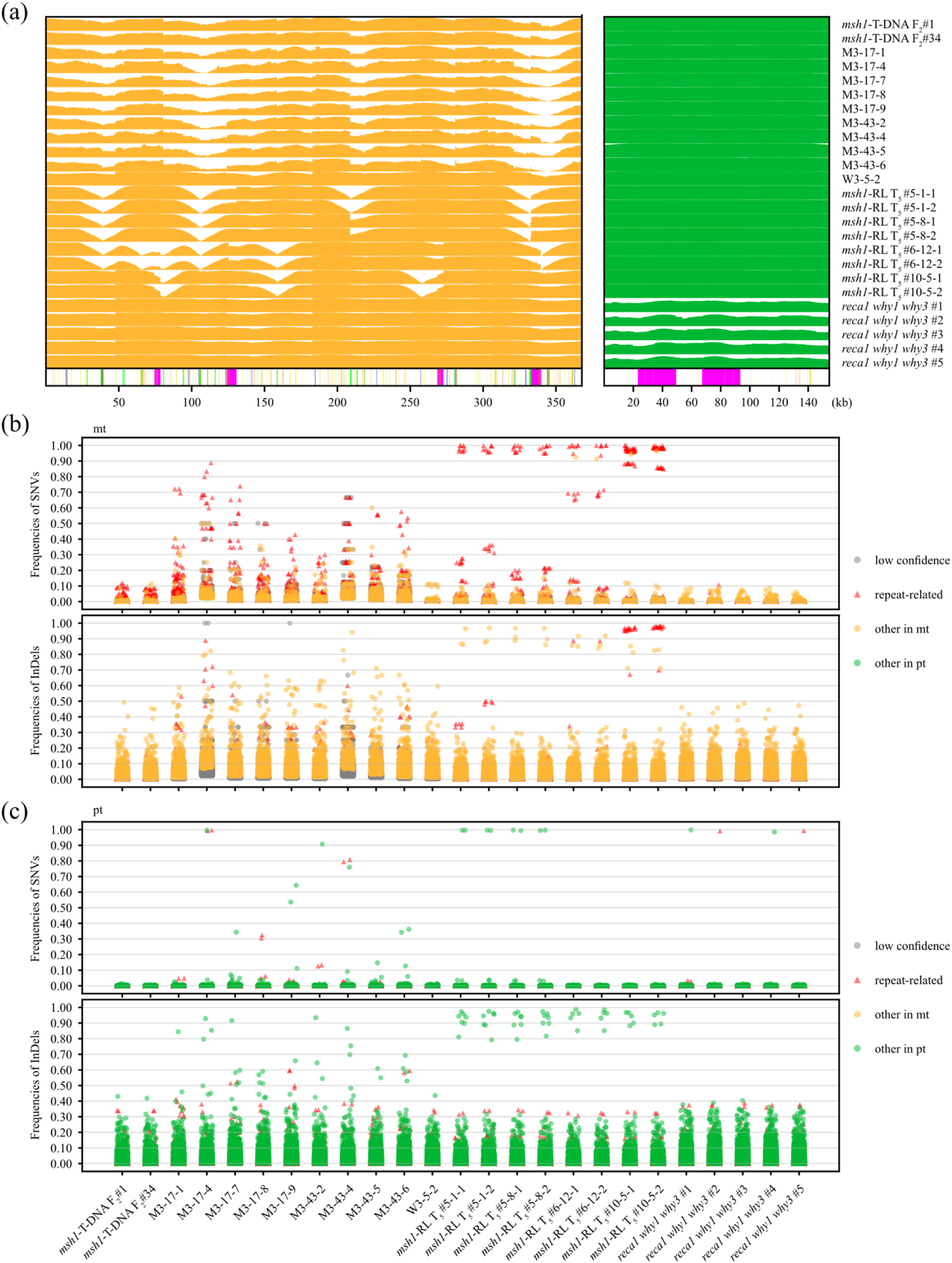
Fixation of small-scale variants in *MSH1* rescue lines and *reca1 why1 why3* triple mutants. (a) Ratios of reference conformations across the mt (orange) and pt (green) genomes in different genetic backgrounds. Repeat annotations (bottom) were shown. (b, c) Frequencies of SNVs and small InDels in (b) mt and (c) pt genomes in different genetic backgrounds.

### Alterations in pt genomes of *recA1 why1 why3* triple mutants

We additionally investigated the consequences of disrupting nuclear-encoding genes with organellar RRR functions by sequencing *recA1 why1 why3* triple mutants, which have a severe leaf variegation phenotype and were previously shown to cause global pt genome instability (Zampini et al., 2015). We found that the dominant pt genome structures were the same as wild type Col-0 even in advanced-generation (self-pollinated for multiple generations) *recA1 why1 why3* triple mutants, although there were genome-wide increases in SV frequencies and rare instances of fixed SNVs (Figure 6c and Figure S22). Collectively, our results describe the frequencies of different types of variants in different genetic backgrounds, providing a quantitative understanding of the raw materials for organellar genome evolution.

## Discussion

In this work, we quantitatively examined the organellar pangenomes using the 149 published HiFi datasets in the model plant *A. thaliana*, which facilitated high-quality assemblies that were validated through a comprehensive estimation of variant frequencies. Combining results from mutants lacking organellar RRR machinery, our analyses integrated both the dominant conformation and dynamic heteroplasmic variants in organellar genomes, providing an unprecedentedly clear view of the complex structural rearrangements of plant mtDNA.

### Flexibility and constraints in the evolution of *A. thaliana* mt and pt genomes

The physical structures of plant mtDNA molecules exist in branched linear or circular forms, which differ significantly from the master circles (Bendich, 1993; Kozik et al., 2019). However, the assembly graphs of wild-type *A. thaliana* accessions, according to our analysis (Figure S1; Zou et al., 2022) and a recent independent study (Xian et al., 2025a), suggested the existence of dominant conformations. The mt and pt genomes exist in heteroplasmic states, which can become increasingly variable in mutants or under stress conditions. Representing different structural forms of mt genomes in assembly graphs would complicate downstream analysis, while still losing some low-frequency forms (He et al., 2023; Bi et al., 2024; Xian et al., 2025b). To accurately determine the master conformation and comprehensively describe variant frequencies, we developed an enhanced version of the hifisr pipeline (Zou et al., 2022) to generate sample-specific variant snapshots with read-level resolution. By comparing the dynamics of organellar genome variants within individual samples and among different *A. thaliana* accessions, we confirmed that the frequencies of structural and small-scale variants varied significantly, both by type and by organelle. Within individuals, large repeats (Large1, Large2, and IRab) and long homopolymers were the most variable regions compared to the master conformations. In Class 3 mt genomes, the number of reads supporting reference conformations far exceeds those of recombined conformations at the 1038-bp Large3 repeat region, suggesting that its recombination was similarly suppressed by organellar RRR machinery as intermediate-sized repeats. For homopolymers of a given size, the average frequency of variants and/or sequencing errors were similar in both mt and pt genomes (Figure 2c). For SNVs, more sites with elevated heteroplasmy frequencies could be observed in mt genomes than in pt genomes (Figure 6b, c). Across accessions, the situation was reversed, where the density of small-scale variants, including homopolymers and SNVs, was more than four-fold lower in mt genomes (Figure 4a; Table S10 and S11). This indicated that the effects on organellar RRR machinery on suppressing heteroplasmy are not directly proportional to divergence across accessions.

Large and intermediate-sized repeats are the main drivers of rapid structural rearrangements during mt genome evolution. Their rearrangements occurred so rapidly that they were often decoupled with other types of variants, similar to the situation observed in centromeres and chromosome arms (Wlodzimierz et al., 2023). Due to the lack of sequence conservation across species, they are particularly well-suited for population-level studies within a species. Unlike pt genomes, mt genomes with more than one pair of large repeats were common (Wynn and Christensen, 2019). In Class 1 mt genomes, two pairs of large repeats combined with a series of intermediate-sized repeats presumably create more structural flexibility and instability (Figure 2a). Loss of genomic fragments in the master conformation was restricted by functional genes. Because the additional sequence around Large1-2 introduced redundancy for genes such as *atp6*, multiple deletion options that resulted in the conversion from Class 1 conformations to Class 2 conformations existed. Among the 56 accessions with Class 2 mt genome conformations, 13 cases were mediated by H-1 and H-2, while 38 were mediated by H-1 and H-3 (Figure 2a; Table S5), a pattern first reported by Christensen (2013) in the comparison between Col-0 and C24. Based on the sequence variation across accession, Large2 repeats seemed more divergent than Large1 (Table S6), but determining whether the emergence of Large2 repeats was earlier than Large1 requires more evidence. These large repeats in mt genomes turnover rapidly, as evidenced by their inconsistent sequences even among closely related species (Figure S8). Our results suggested a possible model for their generation, where longer repeats could form through the elongation or fusion of shorter repeats (Figure 2c), which can be adjacent or distant from each other (Figure S7). In particular, the CPX-4 event in Geg-14 indicated the formation of a pair of 19-kb large repeats through the extension of repeat C (Figure S6c). Although rare in the current set of samples, the generation and loss of large repeats could happen repeatedly over a longer timescale, until intermediate stages became eventually untraceable across species.

These recombination events generate novel structural rearrangements that can potentially affect plant survival and reproduction. For example, the mt genome in Sha accession had one of the two unexplained CPX events. This dramatic change could lead to conflicts between the nuclear and mt genomes, resulting in pollen death and the CMS phenotype (Gobron et al., 2013; Simon et al., 2016; Ricou et al., 2025). Recombination events may also contribute to local adaptation or the expansion of habitats. A recent work using the same HiFi datasets also explored the structures diversity of mt genomes in *A. thaliana* (Xian et al., 2025a). They removed large repeats from the assembly graphs and clustered the contigs based on structural rearrangements, which largely agreed with results of our mt groups. They also used short-read data from hundreds of accessions, attempting to link the loss of Large1 repeats with adaptive advantages (Xian et al., 2025a). However, analysis based on mapping short reads will inevitably lose detailed information of SVs in the master conformation of mt genomes (Theeuwen et al., 2024). Therefore, we urgently need more long-read data and high-quality assemblies to investigate this issue (Alonso-Blanco et al., 2024; The 1001 Genomes Plus Consortium).

### Engineering nuclear-encoded RRR genes to accelerate organellar genome modification

We previously knew that loss of MSH1 caused a global increase in structural and base-level heteroplasmy in organellar genomes. However, results from those studies were primarily derived from pooled samples or merged data from multiple individuals due to limited mt-mapped reads (Arrieta-Montiel et al., 2009; Davila et al., 2011; Wu et al., 2020b; Zou et al., 2022). To make the picture more complete, we examined the F_2_ generation of *msh1* mutants and found lower variant frequencies, suggesting that a prolonged loss is required. We also generated *msh1* rescue lines and observed rearrangements mediated by Large1, Large2, MMJS, D, H, O, and Y. The involvement of MMJS, O, and Y were not observed in the natural accessions we analyzed, possibly due to the limitations of the evolutionary history of the samples. Theoretically, rearrangements mediated by other repeats could be expected if more rescue lines or a broader range natural accessions were examined. In one *msh1* rescue line, the dominant mt genome conformation was not circular with deletion of the redundant sequence near Large1-2, supporting the notion that the multiple deletion pathways were constrained by functional genes. Since our mutants were in the Col-0 background, the most unstable region was centered around Large1-2, which may not be representative of the typical condition for the *A. thaliana* species (Figure 6a and Figure S19). The frequencies of pt SVs in *msh1* mutants were far lower than in *recA1 why1 why3* triple mutants, although the dominant pt genome structures remained unchanged in both backgrounds. Nevertheless, SVs in pt genomes were still observed across *A. thaliana* accessions (Figure 3a and Table S4) and would be more common across species (Wang et al., 2024a).

Organellar genome engineering typically starts from wild-type backgrounds. However, we can unlock the regulation of RRR machinery to increase the types and frequencies of variants, followed by the construction of rescue lines and fixation of the desired variants. This approach mimics natural conditions, where the RRR genes may be regulated by environmental and developmental cues, potentially greatly enhancing the available diversity for organelle genome editing. In conclusion, our research deepens the understanding of the dynamic processes driving organellar genome variation in plants, offering new insights into their structural complexity and evolutionary patterns.

## Methods

### Plant materials, DNA extraction and PacBio HiFi sequencing

To generate the *msh1* rescue lines (*msh1*-RL), we first constructed the vector *MSH1pro-MSH1gDNA-GFP* in the pCambia 1300 backbone. The construct was then transformed into *Agrobacterium tumefaciens* (GV3103) before being introduced into F_3_ families of *msh1* mutants (M3-17 and M3-43 described in Zou et al., 2022), using the floral dip method (Clough and Bent, 1998). Positive T1 transgenic plants were selected on half-strength Murashige and Skoog (½MS) medium containing 1% sucrose, 0.6% agar and 15 mg/L hygromycin B. Plants homozygous for the vector insertion were verified in later generations. Eight T5 individuals (two for each of the four different rescue lines #5-1, #5-8, #6-12, and #10-5) were used for sequencing in this study, i.e., *msh1* (M3-43) / *MSH1pro-MSH1gDNA-GFP* #5-1 and #5-8 derived from the T1 plant #5, *msh1* (M3-17) / *MSH1pro-MSH1gDNA-GFP* #6-12 derived from the T1 plant #6, and *msh1* (M3-17) / *MSH1pro-MSH1gDNA-GFP* #10-5 derived from the T1 plant #10.

To generate the *msh1*-T-DNA lines, we used *msh1* T-DNA mutant (SALK_041951 described in Arrieta-Montiel et al., 2009) as the pollen donor to cross with wild-type Col-0. The F1 plant was self-pollinated, and two different F2 individuals #1 and #34 were used for sequencing in this study. The *reca1 why1 why3* triple mutant was first described in Zampini et al., 2015, and the seeds were provided by Professor Wang Hong-Bin from Guangzhou University of Chinese Medicine.

Plants were grown in a short-day (8-h day / 16-h night) for 6 weeks. Rosettes of each plant individual were collected for DNA extraction and library construction using similar methods described in Zou et al., 2022. Sequencing was performed with the PacBio Sequel II system for *reca1 why1 why3* triple mutants, and the Revio system for *msh1*-T-DNA lines and *msh1* rescue lines.

### The assembly and correction of mt and pt genomes

Our bioinformatics pipeline contains three stages with ten steps (Figure 1a). In the Assembly stage, we first used minimap2 (version 2.26) to extract mt- and pt-mapped reads using the Col-0 mt genome (NC_037304.1) and pt genome (NC_000932.1 plus a 1-bp expansion at position 28,673) as the bait sequence, respectively. We did not use the nuclear sequences to remove NUMT/NUPT signals as in Zou et al. (2022), because the nuclear as well as the organellar genomic variants between accessions could cause unpredictable judgements of primary alignments. To improve the speed of assembling, we randomly sampled 10,000 reads longer than 1 kb (if there were enough reads) and used the correct module of MECAT2 (v20190314; parameters for expected genome sizes are 370 kb for mitochondria and 155 kb for plastid; Xiao et al., 2017) to reduce the amount of the sampled reads. Both the MECAT2-corrected consensus reads, and the sampled reads were assembled using metaFlye (Flye version 2.9.2-b1795; parameters: --meta --pacbio-hifi --extra-params output_gfa_before_rr=1 --genome-size 370K/155K; Kolmogorov et al., 2020), respectively. The assembly graphs before and after automatic repeat resolution by metaFlye were visualized and checked using Bandage (version 0.9.0; Wick et al., 2015). We removed the contigs and links with few support supporting reads and manually resolved the large repeats to get the linear representations of mt and pt genomes (for some accessions, the large repeats in the mt genomes were re-resolved, i.e., the two subgenomic circles were recombined to get the master circle). For mt genomes that showed large divergence from Col-0, we used the linear draft genome of that accession as the bait sequence to perform an additional round of mapping, assembly and repeat resolution, which could ensure the completeness of mt-related information for assembly.

In the Polishing stage, all the linearized genomes were first rotated to make sure that the large repeats were away from the reference boundaries, which was a necessary step for the correct mapping of HiFi reads. For pt genomes, we used 10,000 sampled reads as described above to correct potential errors in these draft assemblies with Flye (version 2.9.2-b1795; parameters: --polish-target). For mt genomes, we used 30,000 mt-mapped reads (if there were enough reads) and remove genome contamination reads with minimap2 (version 2.26; Li et al., 2018) and BamTools (version 2.5.2; parameters: split-reference; Barnett et al., 2011). These contamination reads accounted for approximately two thirds of the input and likely originated from mt genome fragments with sequence similarity to pt genomes. For the Lu-1 accession, we used the rotated mt genome without polishing. To obtain the nuclear genomes of Lu-1 (not provided by Lian et al., 2024) and Col-CEN (incorrect large NUMT as reported by Fields et al., 2022), we performed a de novo assembly using hifiasm (version 0.19.7; parameters: −l 0 -f 0; Cheng et al., 2021), followed by scaffolding the primary contigs (p_ctg) using RagTag (version, 2.1.0; Alonge et al., 2022) base on alignments to Col-PEK (Hou et al., 2022).

In the Validation stage, we first used the same sampled pt-mapped reads and mt-mapped reads (without pt genome contamination) to estimate the heteroplasmy levels of SVs by hifisr (Zou et al., 2022). If the frequency of a given SV had a higher frequency than the accession’s reference conformation, and this SV was not associated with large repeats (> 1 kb), we manually corrected this SV and ran hifisr again. After all erroneous SVs were corrected, we used pt-mapped reads and mt-mapped reads without rearrangements to estimate the frequencies of SNVs and small InDels by Winnowmap2 (version 2.03; Jain et al., 2022), BCFtools (version 1.18; parameters: bcftools mpileup --indels-2.0 -m 1 -Ou; and bcftools call -mv -P 0.99 -Ov; Li et al., 2011) and pysam (version 0.22.0). Then, we manually corrected any erroneous SNVs and InDels and ran the pipeline again. The final variant frequencies were re-calculated using all the reads that fully aligned to the mt and pt genomes. In the last step, we aligned the validated mt and pt genomes to Col-0 by rotation, reverse complementation, or flipping (only for pt genome) using a custom Python script based on BLASTn (version 2.14.1+; Camacho et al., 2009). For variants with frequencies around 50%, we also performed the validation analyses using all the mt- and pt-mapped reads.

### Annotation of mt and pt genomes

To automatically find repeats in organelle genomes, we performed a first round of BLASTn (version 2.14.1+; parameters: -word_size 4; Camacho et al., 2009) search using the mt (or pt) genome for each accession as both query and subject. For each candidate fragment or BLASTn alignment, we removed smaller fragments if they were contained in at least one larger fragment and sorted the remaining fragments based on ascending coordinates. Then we performed two additional rounds of BLASTn (version 2.14.1+; parameters: -word_size 11; Camacho et al., 2009) searches using each sorted fragment in the former round as subject, and this same fragment (to find internal alignments) or the full-length genome (to find multiple alignments) as query. This procedure successfully identified other copies and internal smaller fragments for each fragment and could be applied recursively until no additional internal alignments were found for all fragments. The results of three BLASTn rounds were further processed to remove duplicate alignments and define the copy numbers of repeat groups (size longer than 50 bp). We implemented this repeat annotation algorithm in a custom Python script. We also manually checked the annotations of repeat boundaries and matched the name of repeats in mt genomes with our previous study on Col-0 (Zou et al., 2022). To facilitate the comparison of repeat copies within and between accessions, the conformations and boundaries of repeat copies in other accessions were determined according to BLASTn alignments against the Col-0 reference. Thus, the reference (parent) conformations in some accessions could be the recombinant conformations in Col-0.

For smaller repeats (< 50 bp, also called micro-homologies for the smallest ones), we extracted the 100-bp surrounding sequence for each case of known genomic variants and combined the results of BLASTn (version 2.14.1+; parameters: -word_size 4; Camacho et al., 2009) alignments, naive sequence match, and manual inspection. We did not annotate the complete set of small repeats due their high frequency of random occurrence nature and correspondingly high computational cost. For tandem repeats and partial palindromic repeats, we performed similar analyses and limited the scope to known genomic variants.

To annotate genes encoding proteins, tRNAs, and rRNAs, we performed BLASTn searches using each annotation from Col-0 mt (NC_037304.1) and pt (NC_000932.1) genomes as the subject, and the target mt and pt genomes as queries. For genes with multiple copies, we added suffixes (v1, v2, etc.) to the original gene names. We did not consider ORF genes and RNA editing events in the current analysis.

### Comparative analysis of mt and pt genomes of different accessions

To find the SVs in pt genomes, we performed a pair-wise comparison for each pt genome against the rotated version of Col-0 pt genome using a custom Python script based on BLASTn (version 2.14.1+0; Camacho et al., 2009). The results were manually checked to remove alignments resulted from dual mapping of small fragments. The remaining alignments were counted as true SVs relative to Col-0. For each BLASTn alignment, we analyzed the SNVs and small InDels by Winnowmap2 (version 2.03; Jain et al., 2022) and BCFtools (version 1.18; parameters: bcftools mpileup --indels-2.0 -m 1 -Ou; and bcftools call -mv -P 0.99 -Ov; Li et al., 2011).

To find the SVs in mt genomes, our initial comparisons with Col-0 suggested that rearrangements-mediated by Large1, Large2, B, C, and Q occurred frequently in Class 1 mt genomes, while additional repeats such as H, I, and L were involved in Class 2 and Class 3. Starting from the Col-0 conformation, we flipped the inverted repeats one by one (limited to Large1, Large2, B, C, and Q), which generated a network of 21 possible conformations. We then aligned the mt genome of each accession to these 21 conformations (nodes) and chose the one with the smallest number of BLASTn alignments (dual-mapped fragments were manually checked and removed). We further simplified the mt conformation groups by flipping the large inverted repeats (Large1 and/or Large2), which was similar to the flipping IRab in pt genomes, if their conformations were different from the Col-0 reference. The conformations before and after flipping Large1, Large2 and IRab were both meaningful because of the constitutive recombination mediated by large repeats. Thus, the 140 mt genomes were classified into different conformation groups. Additional SVs were revealed by comparing the mt genomes to the selected accession within each group. To analyze the SNVs and small InDels in mt genomes, we artificially converted their native conformations (corresponding to the majority of heteroplasmies, or master circles) into the Col-0 conformation, and performed analysis using Winnowmap2 (version 2.03; Jain et al., 2022) and BCFtools (version 1.18; parameters: bcftools mpileup --indels-2.0 -m 1 -Ou; and bcftools call -mv -P 0.99 -Ov; Li et al., 2011) as we did for pt genomes. These variants were relative to Col-0 references and were not polarized according to common ancestors, in order to keep sites that were not alignable to outgroup species. The variants within imperfect repeats that mediated rearrangements were analyzed separately in their native conformations, which was important to recover the distribution of true breakpoints.

### Annotation and integration of structural and small-scale variants

We classified the SVs into different types including large InDels, inversion, rearrangement mediated by direct repeats (or micro-homologies) and unalignable complex cases (CPX). We further tried to explain SNVs, small InDels, and SVs by different elementary mutational types according to their related sequence features. We focused on three categories of repetitive sequences: (1) tandem repeats, the repeat units include singletons (homopolymers), dinucleotides and trinucleotides, as well as arbitrary stretches (usually 4 to 6 bases, or more) that could occur at overlapping regions in different accessions; (2) non-tandem repeats (or micro-homologies for very small ones); (3) perfect or partial palindromic repeats. For a given SNV or small InDel variant identified by BCFtools, if its location was not close to other variants, and it could be explained by elementary mutational events mediated by the above sequence features, we called it a simple site. Otherwise, it was identified as a complex site. We manually checked the complex sites to see whether they were related to gene conversion events (exchanges of internal variants between copies of imperfect non-tandem repeats), or micro-inversions of partial palindromic repeats. Based on the comparisons of multiple accession-pairs, we could elucidate some complex sites that were caused by overlapping simple sites if all intermediate mutational steps existed.

To construct the mt and pt pan-genomes, we used the Col-0 references as the backbone sequences and integrated variant calls from all accessions. For simple sites, we kept the Col-0 allele because there were easily identified by BCFtools. The final result consisted of a linear mt pan-genome, a linear pt pan-genome, a few unalignable sequences corresponding to complex presence/absence events CPX-1, CPX-2, and CPX-3.

### Analyzing the consequences of variants for protein coding genes in mt and pt genomes

The effects of SNVs in coding regions were annotated as synonymous, missense, stop gain, or stop loss, while those in intergenic, intron, UTR regions were annotated as non-coding. The effects of two SNVs in the same codon were manually corrected. For small InDels, non-coding, coding border, coding center were assigned. The effects were annotated at the DNA level, and RNA editing events were not considered currently.

### Clustering the accessions using SNVs in pt genomes

To build the pt-SNV tree, we used all the di-allelic simple sites of pt SNVs to calculate the pairwise Hamming distance matrices with the Python package scikit-learn (version 1.5.2) and performed hierarchical clustering using the average linkage method with the Python package scipy (version 1.9.3).

## Data and Code Availability

The raw sequence data for *A. thaliana* accessions (Naish et al., 2021; Hou et al., 2022; Rabanal et al., 2022; Wang et al., 2022; Christenhubz et al., 2023; Kang et al., 2023; Wlodzimierz et al., 2023; Lian et al., 2024), *A. lyrata* (Kolesnikova et al., 2023) and *Capsella bursa-pastoris* (Penin et al., 2024) were downloaded from published studies. The raw sequence data for *msh1*-T-DNA mutants, msh1 rescue lines, and *recA1 why1 why3* triple mutants have been deposited in the Genome Sequence Archive (Chen et al, 2021) in the National Genomics Data Center (CNCB-NGDC Members and Partners, 2025), China National Center for Bioinformation / Beijing Institute of Genomics, Chinese Academy of Sciences (PRJCA035333), which is publicly accessible at https://ngdc.cncb.ac.cn/. The sequences of assemblies and other related information were available at FigShare: https://doi.org/10.6084/m9.figshare.28238345. The updated version of the hifisr pipeline for estimating variant frequences of organellar genomes, including custom scripts referred to in this study, is available at https://github.com/zouyinstein/hifisr.

## Author contributions

ZW designed the research; YZ and YH prepared the plant materials; YZ wrote the computer scripts and performed majority of the bioinformatic analysis; YZ and WZ checked and corrected the assemblies; YZ and YH checked the annotations of organellar genes and repeats; YZ, WZ, DBS, and ZW discussed the results; YZ, DBS and ZW wrote the manuscript; YZ, DBS and ZW revised the manuscript.

## Acknowledgements

This work was supported by grants from the National Natural Science Foundation of China (32170238, 32400191), Basic and Applied Basic Research Foundation of Guangdong Province (2023A1515111029), the Science, Technology and Innovation Commission of Shenzhen Municipality (RCYX20200714114538196), the Chinese Academy of Agricultural Sciences Elite Youth Program (grant 110243160001007), the Guangdong Pearl River Talent Program (2021QN02N792) and US National Institutes of Health (R35GM148134).

## List of Supplementary Figures and Tables

**Figure S1.** Bandage view of metaFlye organellar genome assembly graphs.

**Figure S2.** Rotation of the reference genome ensures correct mapping of HiFi reads.

**Figure S3.** Examples of structural variants with higher frequencies in *A. thaliana* accessions.

**Figure S4.** Samples with different dominant lengths at long homopolymers of the same accession.

**Figure S5.** An example with high NUMT noise in Lu-1 accession.

**Figure S6.** Revealing the complex (CPX) events by flipping the reference genome.

**Figure S7.** Examples of repeat elongation and fusion events in mt genomes.

**Figure S8.** The mt genomes became highly rearranged across species.

**Figure S9.** Most of the small-scale variants in mt and pt genomes were limited to a small number of samples.

**Figure S10.** Examples of closely located small-scale variants explained by gene conversion events.

**Figure S11.** Examples of closely located small-scale variants explained by micro-inversion events in mt genomes.

**Figure S12.** Examples of closely located small-scale variants explained by micro-inversion events in pt genomes.

**Figure S13.** An example of closely located small-scale variants explained by repeat extension.

**Figure S14.** Some mutations were better explained when comparing multiple accessions.

**Figure S15.** Geographic locations of the 149 samples used in this study.

**Figure S16.** Structures of MSH1 dimers binding to double-stranded DNA with a mismatch.

**Figure S17.** Coverages and SV frequencies in the mt genomes of *msh1* rescue lines.

**Figure S18.** Coverages and SV frequencies in the pt genomes of *msh1* rescue lines.

**Figure S19.** Coverages and SV frequencies in the mt genomes of *msh1* mutants.

**Figure S20.** Frequencies of SVs and small-scale variants were affected by the giant NUMT on Chr2 of Col-0.

**Figure S21.** Coverages and SV frequencies in the pt genomes of *msh1* mutants.

**Figure S22.** Coverages and SV frequencies in the mt and pt genomes of *recA1 why1 why3* triple mutants.

**Table S1**. Statistics of HiFi reads mapped to the mt and pt genomes in 149 samples of *A. thaliana*.

**Table S2**. Samples with different reference sequences for the same accession.

**Table S3**. BLASTn results between the NUMT (Chr2_mt_g9) and mt genome in Lu-1.

**Table S4**. Structural rearrangements in pt genomes of *A. thaliana* accessions.

**Table S5**. Structural rearrangements in mt genomes of *A. thaliana* accessions.

**Table S6**. Small-scale variants within Large1, Large2, IRab in 149 samples.

**Table S7**. Small-scale variants within intermediate-sized repeats that mediated mt genome rearrangements in 149 samples.

**Table S8**. BLASTn results illustrating complex rearrangement events in mt genomes

**Table S9**. BLASTn results between the mt genomes of *A. thaliana* and related outgroup species.

**Table S10**. Small-scale variants in 149 samples compared with Col-0 reference mt genome.

**Table S11**. Small-scale variants in 149 samples compared with Col-0 reference pt genome.

**Table S12**. Effects of SNVs in 149 samples of *A. thaliana*.

**Table S13**. Effects of small InDels in 149 samples of *A. thaliana*.

**Table S14**. Geological information of the 149 samples of *A. thaliana*.

**Table S15**. BLASTn results illustrating mitochondrial structural rearrangements in *msh1* rescue lines.

**Table S16**. Small-scale variants in *msh1* rescue lines compared with Col-0 reference mt genome.

**Table S17**. Small-scale variants in *msh1* rescue lines compared with Col-0 reference pt genome.

## References

Abdelnoor RV, Yule R, Elo A, Christensen AC, Meyer-Gauen G, Mackenzie SA. Substoichiometric shifting in the plant mitochondrial genome is influenced by a gene homologous to MutS. Proc Natl Acad Sci U S A 2003, 100(10): 5968–5973.

Abramson J, Adler J, Dunger J, Evans R, Green T, Pritzel A, et al. Accurate structure prediction of biomolecular interactions with AlphaFold 3. Nature 2024, 630(8016): 493–500.

Alonge M, Lebeigle L, Kirsche M, Jenike K, Ou S, Aganezov S, et al. Automated assembly scaffolding using RagTag elevates a new tomato system for high-throughput genome editing. Genome Biol 2022, 23(1): 258.

Alonso-Blanco CC, Ashkenazy H, Baduel P, Bao Z, Becker C, Caillieux E, et al. The 1001G+ project: A curated collection of Arabidopsis thaliana long-read genome assemblies to advance plant research. bioRxiv 2024.

Arimura SI, Ayabe H, Sugaya H, Okuno M, Tamura Y, Tsuruta Y, et al. Targeted gene disruption of *ATP synthases 6-1* and *6-2* in the mitochondrial genome of *Arabidopsis thaliana* by mitoTALENs. Plant J 2020, 104(6): 1459–1471.

Arrieta-Montiel MP, Shedge V, Davila J, Christensen AC, Mackenzie SA. Diversity of the *Arabidopsis* mitochondrial genome occurs via nuclear-controlled recombination activity. Genetics 2009, 183(4): 1261–1268.

Barnett DW, Garrison EK, Quinlan AR, Stromberg MP, Marth GT. BamTools: a C++ API and toolkit for analyzing and managing BAM files. Bioinformatics 2011, 27(12): 1691–1692.

Bendich AJ. Reaching for the ring: the study of mitochondrial genome structure. Curr Genet 1993, 24(4): 279–290.

Bi C, Shen F, Han F, Qu Y, Hou J, Xu K, et al. PMAT: an efficient plant mitogenome assembly toolkit using low-coverage HiFi sequencing data. Hortic Res 2024, 11(3): uhae023.

Broz AK, Keene A, Fernandes Gyorfy M, Hodous M, Johnston IG, Sloan DB. Sorting of mitochondrial and plastid heteroplasmy in *Arabidopsis* is extremely rapid and depends on MSH1 activity. Proc Natl Acad Sci U S A 2022, 119(34): e2206973119.

Broz AK, Sloan DB, Johnston IG. Stochastic organelle genome segregation through *Arabidopsis* development and reproduction. New Phytol 2024, 241(2): 896–910.

Camacho C, Coulouris G, Avagyan V, Ma N, Papadopoulos J, Bealer K, et al. BLAST+: architecture and applications. BMC Bioinformatics 2009, 10: 421.

Chen L, Liu YG. Male sterility and fertility restoration in crops. Annu Rev Plant Biol 2014, 65: 579–606.

Chen T, Chen X, Zhang S, Zhu J, Tang B, Wang A, et al. The genome sequence archive family: toward explosive data growth and diverse data types. genomics proteomics bioinformatics 2021, 19(4): 578–583.

CNCB-NGDC Members and Partners. Database resources of the National Genomics Data Center, China National Center for Bioinformation in 2025. Nucleic Acids Res 2025, 53(D1): D30–D44.

Cheng H, Concepcion GT, Feng X, Zhang H, Li H. Haplotype-resolved de novo assembly using phased assembly graphs with hifiasm. Nat Methods 2021, 18(2): 170–175.

Christenhusz MJM, Twyford AD, Hudson A. The genome sequence of thale cress, Arabidopsis thaliana (Heynh., 1842). Wellcome Open Research 2023, 8.

Christensen AC. Plant mitochondrial genome evolution can be explained by DNA repair mechanisms. Genome Biol Evol 2013, 5(6): 1079–1086.

Clough SJ, Bent AF. Floral dip: a simplified method for Agrobacterium-mediated transformation of *Arabidopsis thaliana*. Plant J 1998, 16(6): 735–743.

Davila JI, Arrieta-Montiel MP, Wamboldt Y, Cao J, Hagmann J, Shedge V, et al. Double-strand break repair processes drive evolution of the mitochondrial genome in *Arabidopsis*. BMC Biol 2011, 9: 64.

Fields PD, Waneka G, Naish M, Schatz MC, Henderson IR, Sloan DB. Complete sequence of a 641-kb insertion of mitochondrial DNA in the *Arabidopsis thaliana* nuclear genome. Genome Biol Evol 2022, 14(5).

Forner J, Kleinschmidt D, Meyer EH, Gremmels J, Morbitzer R, Lahaye T, et al. Targeted knockout of a conserved plant mitochondrial gene by genome editing. Nat Plants 2023, 9(11): 1818–1831.

Gobron N, Waszczak C, Simon M, Hiard S, Boivin S, Charif D, et al. A cryptic cytoplasmic male sterility unveils a possible gynodioecious past for *Arabidopsis thaliana*. PLoS One 2013, 8(4): e62450.

Golin S, Negroni YL, Bennewitz B, Klosgen RB, Mulisch M, La Rocca N, et al. WHIRLY2 plays a key role in mitochondria morphology, dynamics, and functionality in *Arabidopsis thaliana*. Plant Direct 2020, 4(5): e00229.

Gualberto JM, Newton KJ. Plant mitochondrial genomes: dynamics and mechanisms of mutation. Annu Rev Plant Biol 2017, 68: 225–252.

Gupta R, Kanai M, Durham TJ, Tsuo K, McCoy JG, Kotrys AV, et al. Nuclear genetic control of mtDNA copy number and heteroplasmy in humans. Nature 2023, 620(7975): 839–848.

He W, Xiang K, Chen C, Wang J, Wu Z. Master graph: an essential integrated assembly model for the plant mitogenome based on a graph-based framework. Briefings in Bioinformatics 2023, 24(1).

Hou X, Wang D, Cheng Z, Wang Y, Jiao Y. A near-complete assembly of an *Arabidopsis thaliana* genome. Mol Plant 2022, 15(8): 1247–1250.

Jain C, Rhie A, Hansen NF, Koren S, Phillippy AM. Long-read mapping to repetitive reference sequences using Winnowmap2. Nat Methods 2022, 19(6): 705–710.

Kang M, Wu H, Liu H, Liu W, Zhu M, Han Y, et al. The pan-genome and local adaptation of *Arabidopsis thaliana*. Nat Commun 2023, 14(1): 6259.

Kolesnikova UK, Scott AD, Van de Velde JD, Burns R, Tikhomirov NP, Pfordt U, et al. Transition to self-compatibility associated with dominant S-allele in a diploid Siberian progenitor of allotetraploid *Arabidopsis kamchatica* revealed by *Arabidopsis lyrata* genomes. Mol Biol Evol 2023, 40(7).

Kolmogorov M, Bickhart DM, Behsaz B, Gurevich A, Rayko M, Shin SB, et al. metaFlye: scalable long-read metagenome assembly using repeat graphs. Nat Methods 2020, 17(11): 1103–1110.

Kozik A, Rowan BA, Lavelle D, Berke L, Schranz ME, Michelmore RW, et al. The alternative reality of plant mitochondrial DNA: One ring does not rule them all. PLoS Genet 2019, 15(8): e1008373.

Li H. A statistical framework for SNP calling, mutation discovery, association mapping and population genetical parameter estimation from sequencing data. Bioinformatics 2011, 27(21): 2987–2993.

Li H. Minimap2: pairwise alignment for nucleotide sequences. Bioinformatics 2018, 34(18): 3094–3100.

Lian Q, Hüttel B, Walkemeier B, Mayjonade B, Lopez-Roques C, Gil L, et al. A pan-genome of 69 Arabidopsis thaliana accessions reveals a conserved genome structure throughout the global species range. Nat Genet 2024.

Ma X, Fan J, Wu Y, Zhao S, Zheng X, Sun C, et al. Whole-genome *de novo* assemblies reveal extensive structural variations and dynamic organelle-to-nucleus DNA transfers in African and Asian rice. Plant J 2020, 104(3): 596–612.

Mackenzie SA, Kundariya H. Organellar protein multi-functionality and phenotypic plasticity in plants. Philos Trans R Soc Lond B Biol Sci 2020, 375(1790): 20190182.

Malinova I, Zupok A, Massouh A, Schottler MA, Meyer EH, Yaneva-Roder L, et al. Correction of frameshift mutations in the *atpB* gene by translational recoding in chloroplasts of *Oenothera* and tobacco. Plant Cell 2021, 33(5): 1682–1705.

Martinez-Zapater JM, Gil P, Capel J, Somerville CR. Mutations at the *Arabidopsis CHM* locus promote rearrangements of the mitochondrial genome. Plant Cell 1992, 4(8): 889–899.

Massouh A, Schubert J, Yaneva-Roder L, Ulbricht-Jones ES, Zupok A, Johnson MT, et al. Spontaneous chloroplast mutants mostly occur by replication slippage and show a biased pattern in the plastome of *Oenothera*. Plant Cell 2016, 28(4): 911–929.

Naish M, Alonge M, Wlodzimierz P, Tock AJ, Abramson BW, Schmucker A, et al. The genetic and epigenetic landscape of the *Arabidopsis* centromeres. Science 2021, 374(6569): eabi7489.

Negroni YL, Doro I, Tamborrino A, Luzzi I, Fortunato S, Hensel G, et al. The *Arabidopsis* mitochondrial nucleoid-associated protein WHIRLY2 is required for a proper response to salt stress. Plant Cell Physiol 2024, 65(4): 576–589.

Park HS, Jeon JH, Cho W, Lee Y, Park JY, Kim J, et al. High-throughput discovery of plastid genes causing albino phenotypes in ornamental chimeric plants. Hortic Res 2023, 10(1): uhac246.

Penin AA, Kasianov AS, Klepikova AV, Omelchenko DO, Makarenko MS, Logacheva MD. Origin and diversity of *Capsella bursa-pastoris* from the genomic point of view. BMC Biol 2024, 22(1): 52.

Rabanal FA, Graff M, Lanz C, Fritschi K, Llaca V, Lang M, et al. Pushing the limits of HiFi assemblies reveals centromere diversity between two *Arabidopsis thaliana* genomes. Nucleic Acids Res 2022, 50(21): 12309–12327.

Ricou A, Simon M, Duflos R, Azzopardi M, Roux F, Budar F, et al. Identification of novel genes responsible for a pollen killer present in local natural populations of *Arabidopsis thaliana*. PLoS Genet 2025, 21(1): e1011451.

Sakamoto W, Kondo H, Murata M, Motoyoshi F. Altered mitochondrial gene expression in a maternal distorted leaf mutant of *Arabidopsis* induced by chloroplast mutator. Plant Cell 1996, 8(8): 1377–1390.

Shedge V, Arrieta-Montiel M, Christensen AC, Mackenzie SA. Plant mitochondrial recombination surveillance requires unusual *RecA* and *MutS* homologs. Plant Cell 2007, 19(4): 1251–1264.

Simon M, Durand S, Pluta N, Gobron N, Botran L, Ricou A, et al. Genomic conflicts that cause pollen mortality and raise reproductive barriers in *Arabidopsis thaliana*. Genetics 2016, 203(3): 1353–1367.

Sloan DB, Broz AK, Kuster SA, Muthye V, Penafiel-Ayala A, Marron JR, et al. Expansion of the *MutS* gene family in plants. Plant Cell 2024.

Takatsuka A, Kazama T, Arimura SI, Toriyama K. TALEN-mediated depletion of the mitochondrial gene *orf312* proves that it is a Tadukan-type cytoplasmic male sterility-causative gene in rice. Plant J 2022, 110(4): 994–1004.

Theeuwen T, Wijfjes RY, Dorussen D, Lawson AW, Lind J, Jin K, et al. Species-wide inventory of *Arabidopsis thaliana* organellar variation reveals ample phenotypic variation for photosynthetic performance. Proc Natl Acad Sci U S A 2024, 121(49): e2414024121.

Virdi KS, Wamboldt Y, Kundariya H, Laurie JD, Keren I, Kumar KRS, et al. MSH1 Is a plant organellar DNA binding and thylakoid protein under precise spatial regulation to alter development. Mol Plant 2016, 9(2): 245–260.

Wang B, Yang X, Jia Y, Xu Y, Jia P, Dang N, et al. High-quality *Arabidopsis thaliana* genome assembly with Nanopore and HiFi long reads. Genomics Proteomics Bioinformatics 2022, 20(1): 4–13.

Wang J, Kan S, Kong J, Nie L, Fan W, Ren Y, et al. Accumulation of large lineage-specific repeats coincides with sequence acceleration and structural rearrangement in *Plantago* plastomes. Genome Biol Evol 2024a, 16(8).

Wang J, Kan S, Liao X, Zhou J, Tembrock LR, Daniell H, et al. Plant organellar genomes: much done, much more to do. Trends Plant Sci 2024b, 29(7): 754–769.

Wick RR, Schultz MB, Zobel J, Holt KE. Bandage: interactive visualization of *de novo* genome assemblies. Bioinformatics 2015, 31(20): 3350–3352.

Wlodzimierz P, Rabanal FA, Burns R, Naish M, Primetis E, Scott A, et al. Cycles of satellite and transposon evolution in *Arabidopsis* centromeres. Nature 2023, 618(7965): 557–565.

Wu Z, Waneka G, Sloan DB. The tempo and mode of angiosperm mitochondrial genome divergence inferred from intraspecific variation in *Arabidopsis thaliana*. G3 (Bethesda) 2020a, 10(3): 1077–1086.

Wu Z, Waneka G, Broz AK, King CR and Sloan DB. *MSH1* is required for maintenance of the low mutation rates in plant mitochondrial and plastid genomes. Proc Natl Acad Sci U S A 2020b, 117, 16448–16455.

Wynn EL, Christensen AC. Repeats of unusual size in plant mitochondrial genomes: identification, incidence and evolution. G3 (Bethesda) 2019, 9(2): 549–559.

Xian W, Bao Z, Vorbrugg S, Tao Y, Movilli A, Bezrukov I, et al. The structure of mitochondrial genomes is associated with geography in Arabidopsis thaliana. bioRxiv 2025a.

Xian W, Bezrukov I, Bao Z, Vorbrugg S, Gautam A, Weigel D. TIPPo: A user-friendly tool for *de novo* assembly of organellar genomes with high-fidelity data. Mol Biol Evol 2025b, 42(1).

Xiao CL, Chen Y, Xie SQ, Chen KN, Wang Y, Han Y, et al. MECAT: fast mapping, error correction, and de novo assembly for single-molecule sequencing reads. Nat Methods 2017, 14(11): 1072–1074.

Xu YZ, Arrieta-Montiel MP, Virdi KS, de Paula WB, Widhalm JR, Basset GJ, et al. MutS HOMOLOG1 is a nucleoid protein that alters mitochondrial and plastid properties and plant response to high light. Plant Cell 2011, 23(9): 3428–3441.

Zampini E, Lepage E, Tremblay-Belzile S, Truche S, Brisson N. Organelle DNA rearrangement mapping reveals U-turn-like inversions as a major source of genomic instability in *Arabidopsis* and humans. Genome Res 2015, 25(5): 645–654.

Zhong Y, Okuno M, Tsutsumi N, Arimura S-i. Mitochondrial DNA and the 641kb nuclear-mitochondrial DNA in *Arabidopsis* can be separated by their CpG methylation levels. bioRxiv 2024.

Zhou J, Nie L, Zhang S, Mao H, Arimura SI, Jin S, et al. Mitochondrial genome editing of *WA352* via mitoTALENs restore fertility in cytoplasmic male sterile rice. Plant Biotechnol J 2024, 22(7): 1960–1962.

Zou Y, Zhu W, Sloan DB, Wu Z. Long-read sequencing characterizes mitochondrial and plastid genome variants in *Arabidopsis msh1* mutants. Plant J 2022, 112(3): 738–755.

