## Supplementary Figures for "The evolutionary dynamics of organellar pan-genomes in *Arabidopsis thaliana*"

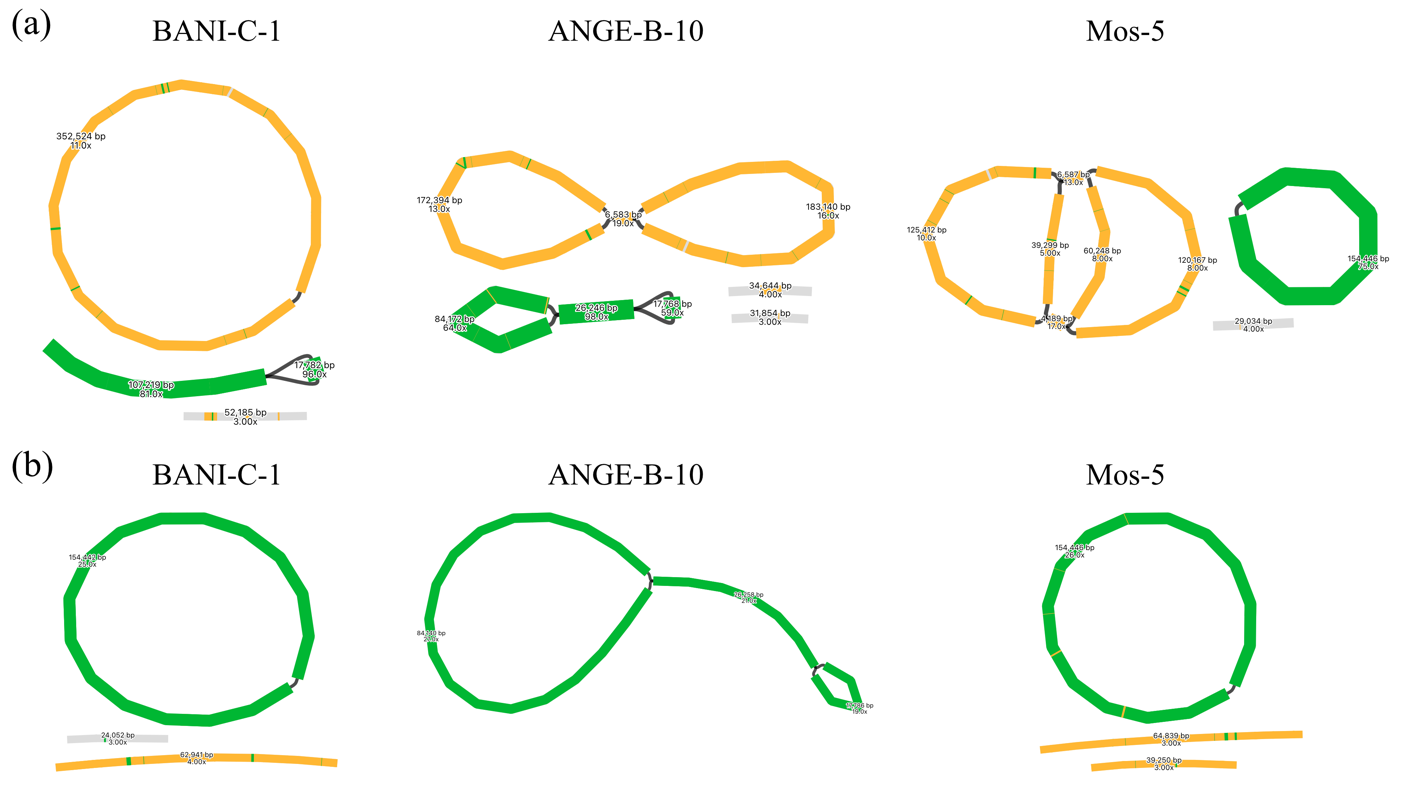

**Figure S1**. Bandage view of metaFlye organellar genome assembly graphs. (a) Assembly graphs of mt genomes in representative accessions with zero (left), one (middle), or two (right) unresolved large repeats. (b) Assembly graphs of corresponding pt genomes. Colors of contigs indicate the BLASTn similarity for mt (orange) and pt (green) references.

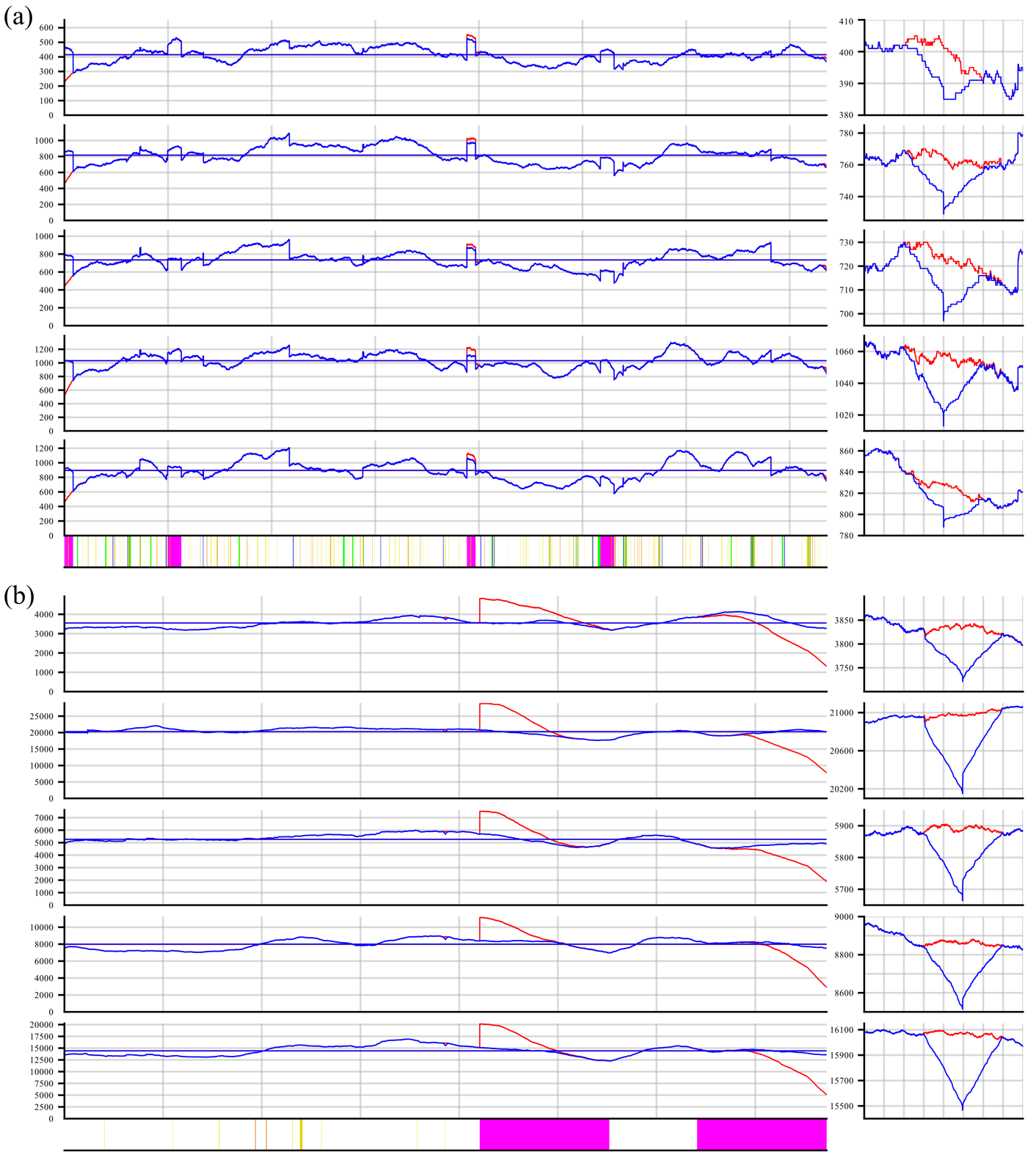

**Figure S2**. Rotation of the reference genome ensures correct mapping of HiFi reads. (a, b) Coverages of five Col-0 samples against the original (red) and rotated (blue) reference (a) mt and (b) pt genomes. The drop in coverage at the new boundaries of the rotated references were shown on the right. Repeat annotations (bottom) were shown.

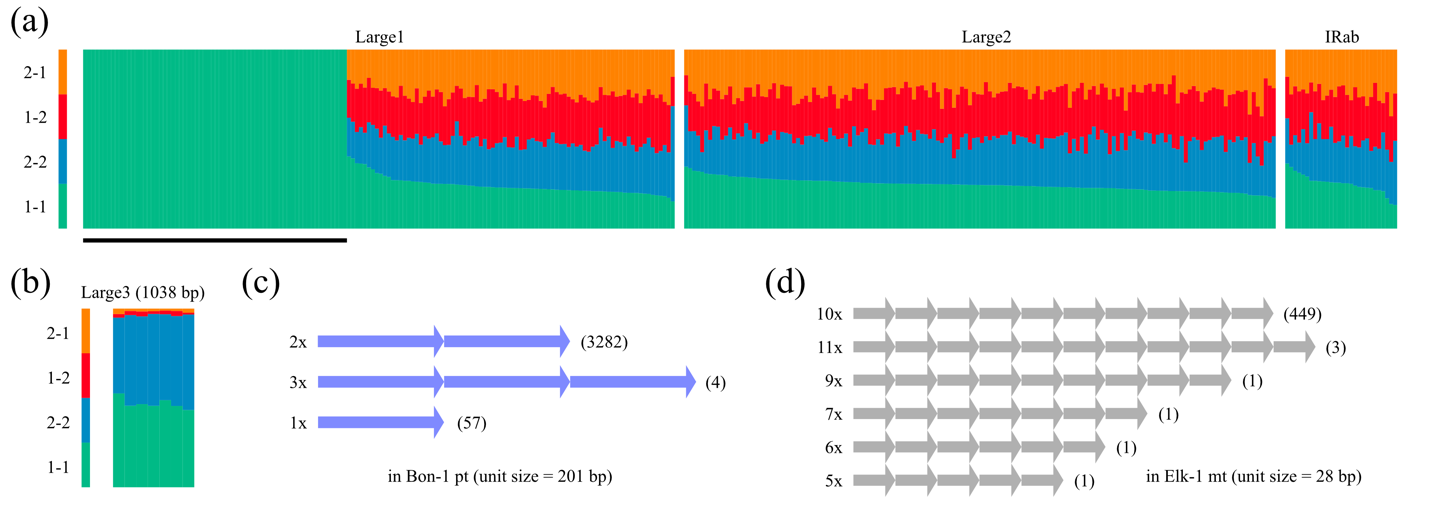

**Figure S3**. Examples of structural variants with higher frequencies in *A. thaliana* accessions. (a) Stacked bar plots show almost equal proportions of reads supporting the reference and recombined conformations at repeats Large1, Large2 and IRab in different accessions. Samples with less than 50 reads supporting these four conformations were removed. 1-1, 2-2 indicate the conformations observed in the reference genome of Col-0, while 1-2 and 2-1 indicate the recombined conformations relative to Col-0 reference. (b, c, d) Examples with slightly increased SV frequencies at (b) Large3 in seven accessions, and tandem repeats with (c) a long unit length (201-bp unit length in Bon-1 pt genome) or (d) a high copy number (ten-copy of a 28-bp unit in Elk-1 mt genome). Numbers in parentheses indicate the number of reads supporting the copy number of tandem repeats. Note that the copy number in the reference genome is supported by most reads.

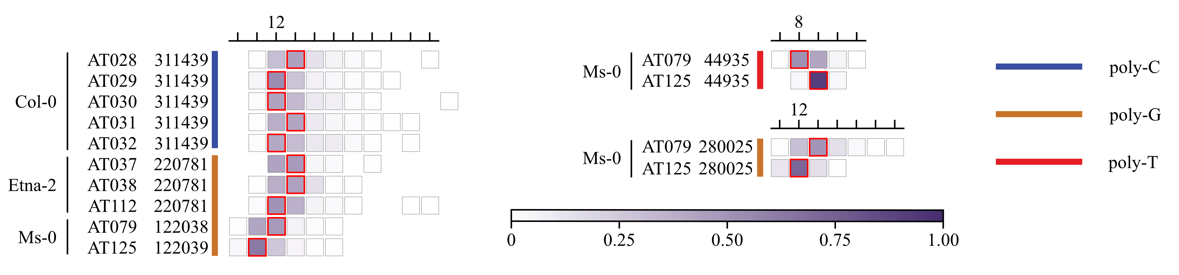

**Figure S4**. Samples with different dominant lengths at long homopolymers of the same accession. The boxes with red borders indicate the most frequent length in the corresponding accession. Heatmaps of allele frequencies in different samples were aligned by the absolute homopolymer length, not by the derivation from the reference length. Position 311439 in Col-0, position 220281 in Etna-2, and positions 122038 and 122039 in Ms-0 were the same homopolymer based on sequence homology of flanking regions.

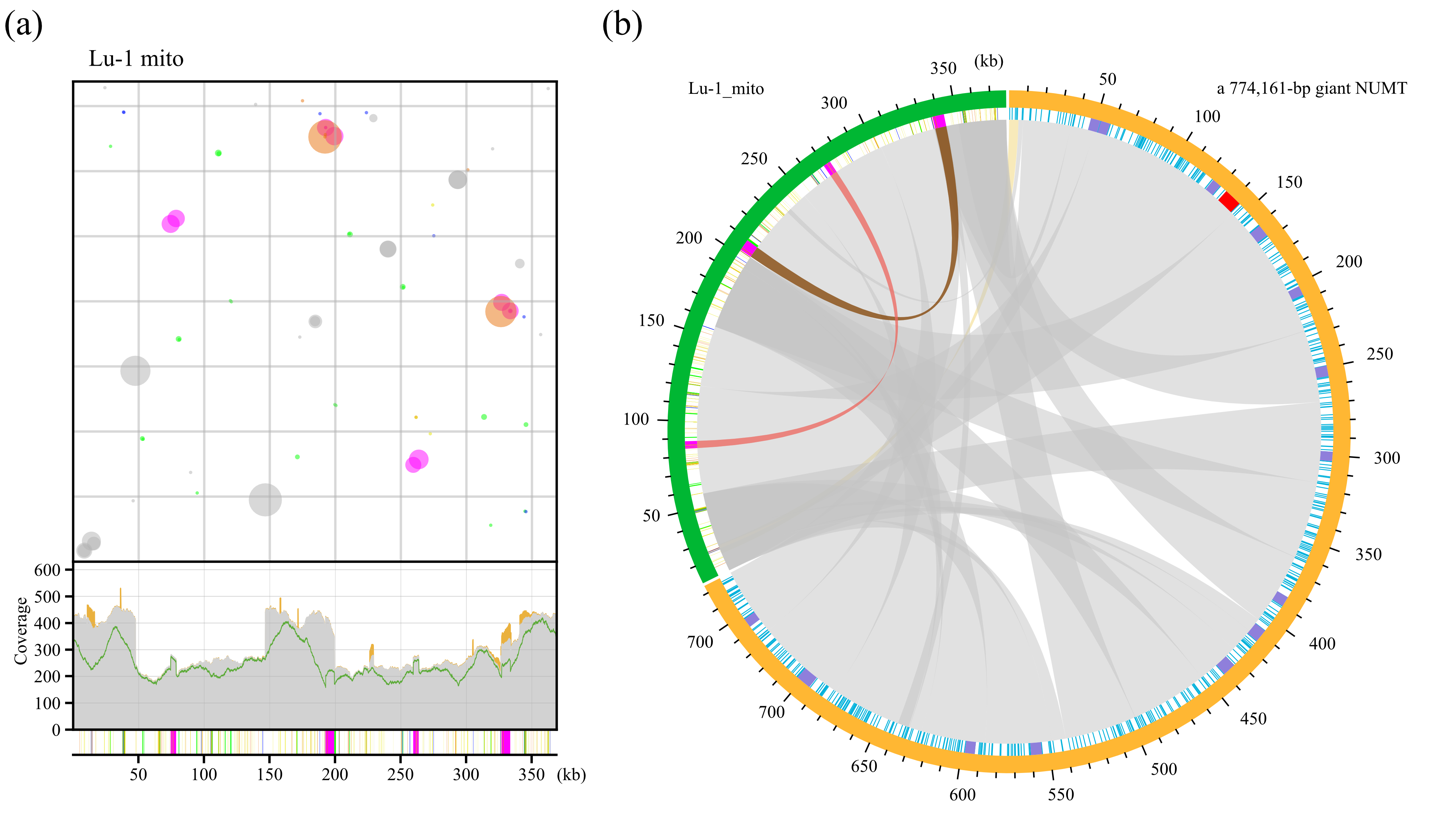

**Figure S5**. An example with high NUMT noise in Lu-1 accession. (a) Frequencies (top) of one-rearrangement reads and coverages (bottom) of HiFi reads were strongly influenced by NUMT-derived signals. In the bubble plot, colors represent categories of alignment overlap length (AO length) as in Zou et al. (2022). In the coverage plot, grey areas indicate fully aligned reads, orange areas indicate partially aligned reads, and the green line indicate reads with no rearrangements plus one-rearrangement reads mediated by Large1 and Large2. (b) Circos plot shows the SVs and small-scale variants between the mt genome (green arc) and the NUMT (orange arc). The track inside repeat annotations in the mt genome and the small-scale variants in the NUMT. The small-scale variants were shown as blue lines, with long identical regions indicated by purple (≥ 5 kb) and red (≥ 10 kb) colors. The chords show direct (grey) and inverted (yellow) syntenic regions based on BLASTn alignments, and large repeats in direct (brown) and inverted (red) orientation are displayed as annotations.

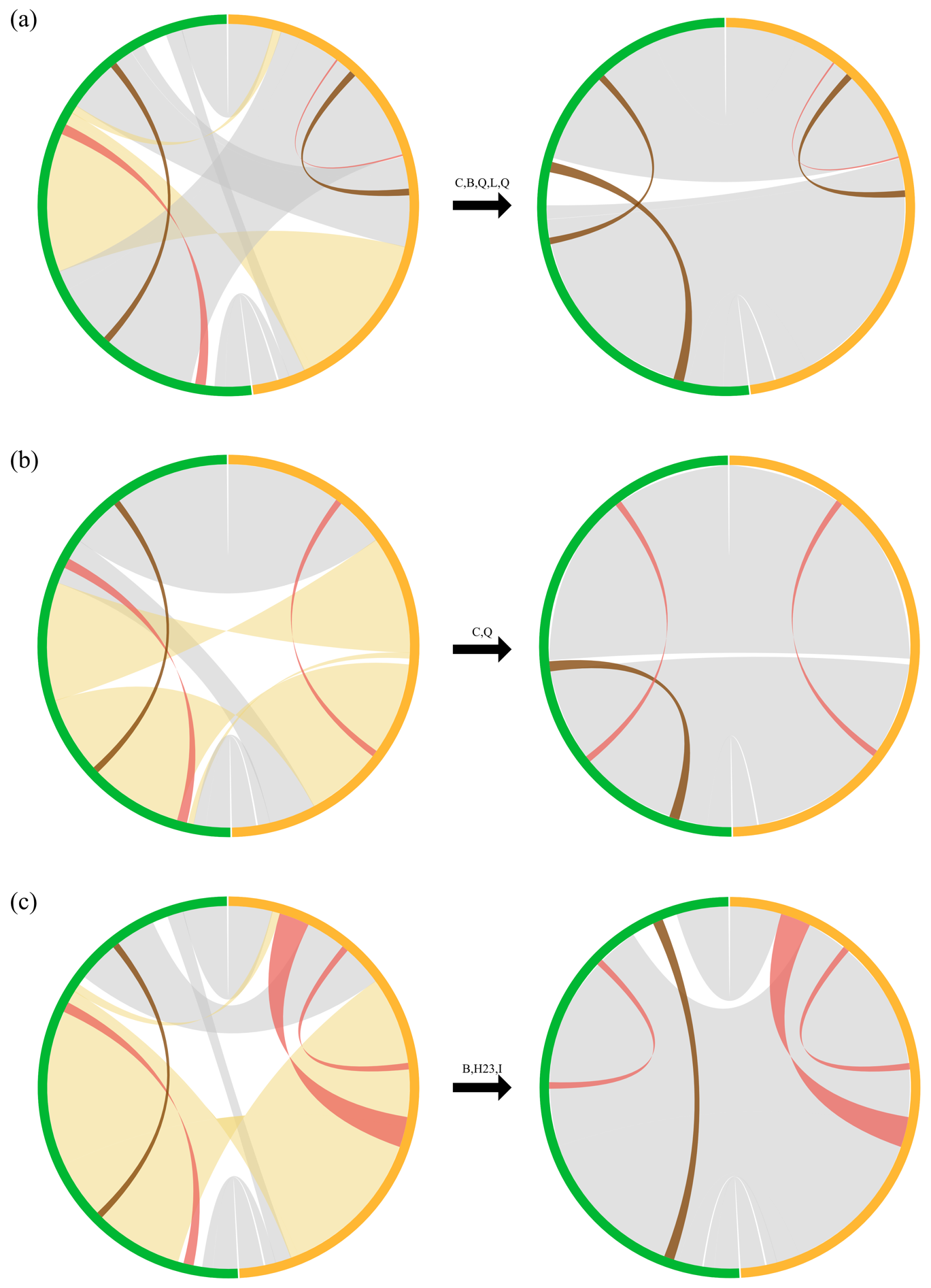

**Figure S6**. Revealing the complex (CPX) events by flipping the reference genome. (a, b, c) Circos plots show the syntenic relationship between the mt genomes with (a) CPX-2, (b) CPX-3, and (c) CPX-4 events (orange arc) and the mt genomes of the original (left) and flipped (right) Col-0 (green arc). The chords show direct (grey) and inverted (yellow) syntenic regions based on BLASTn alignments, and large repeats in direct (brown) and inverted (red) orientation are displayed as annotations.

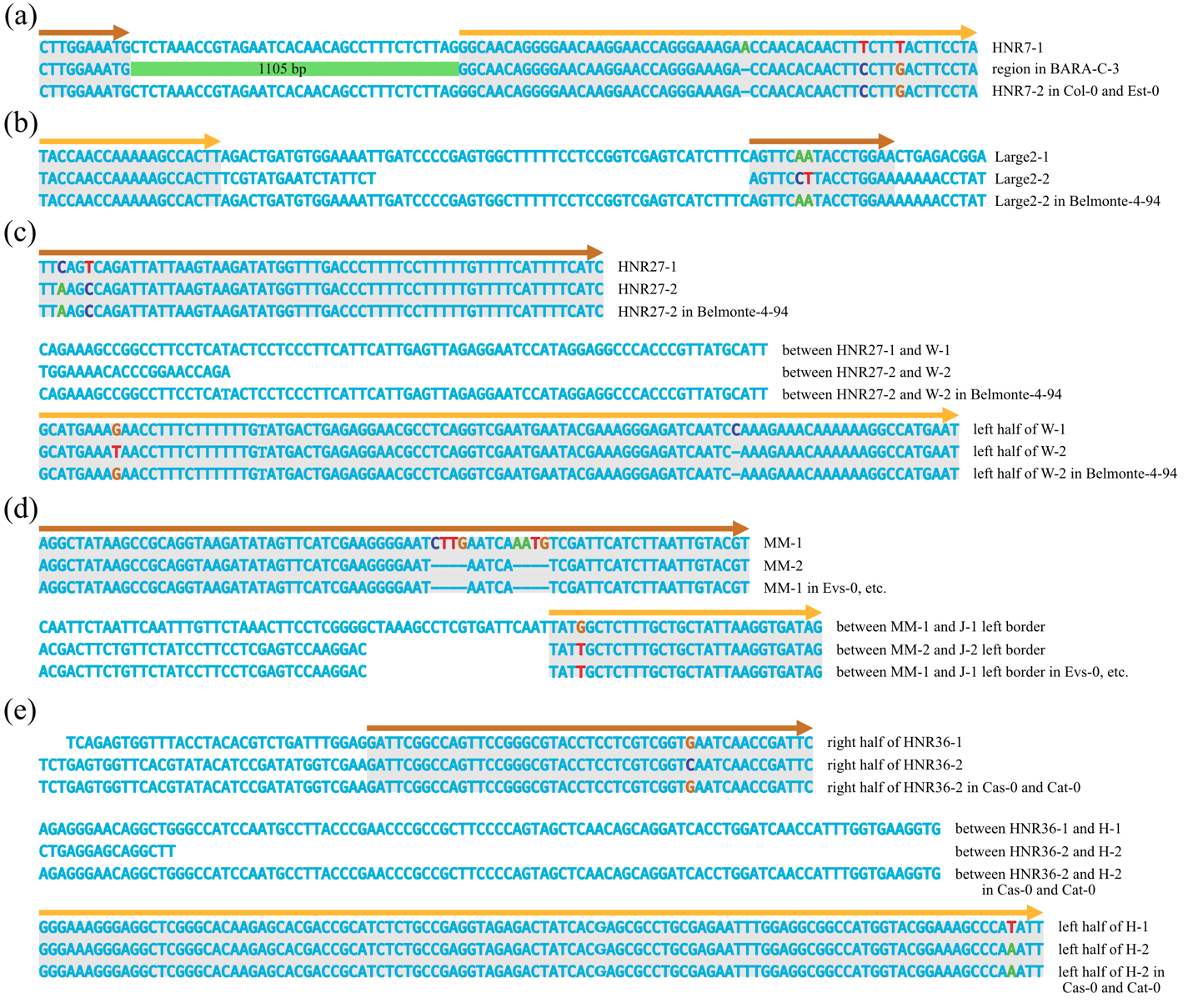

**Figure S7**. Examples of repeat elongation and fusion events in mt genomes. (a, b) Some complex structural rearrangements such as (a) CPX-1 and (b) CPX-5 could be explained by repeat elongation and fusion events. (a, b, c) Additional repeat fusion events not identified as structural variants by BLASTn were also observed.

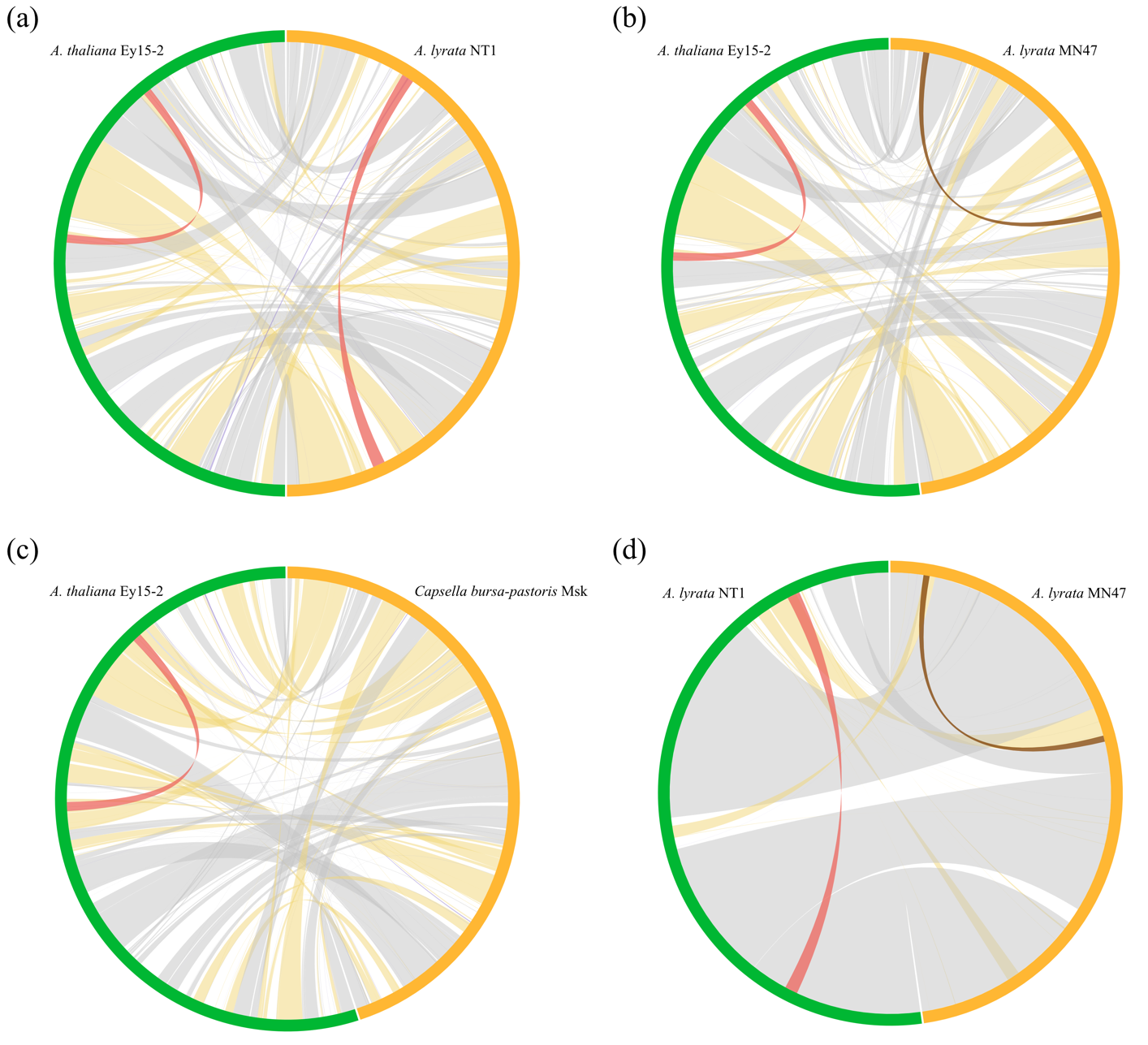

**Figure S8**. The mt genomes became highly rearranged across species. (a-d) Circos plots show the interspecific syntenic relationship between *A. thaliana* Ey15-2 and (a) *A. lyrata* NT1, (b) *A. lyrata* NT1, (c) *Capsella bursa-pastoris* Msk, and the intraspecific syntenic relationship between (d) *A. lyrata* NT1 and MN47. The chords show direct (grey) and inverted (yellow) syntenic regions based on BLASTn alignments, and large repeats in direct (brown) and inverted (red) orientation are displayed as annotations.

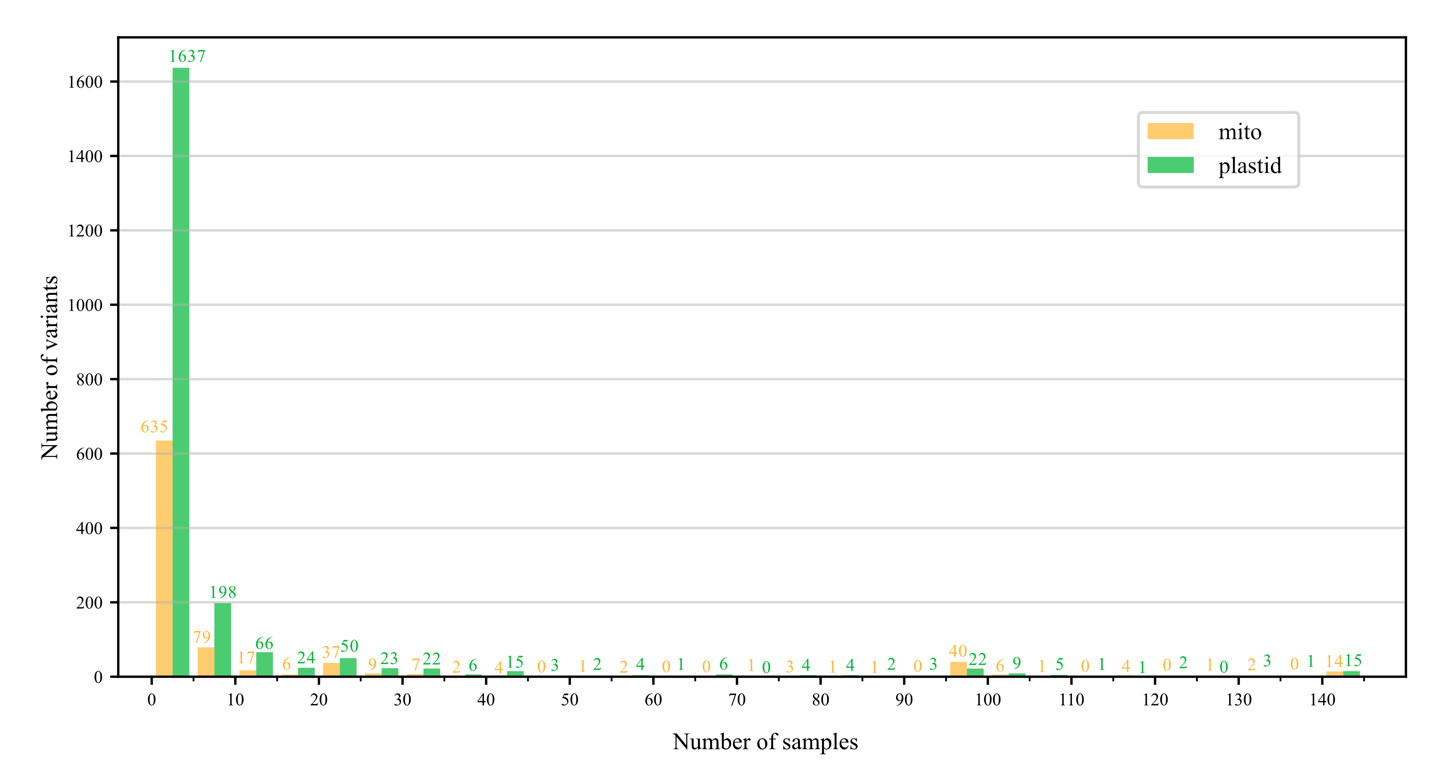

**Figure S9**. Most of the small-scale variants in mt and pt genomes were limited to a small number of samples. Bar plots show the number of variants shared by different sample counts, with the values summed in increments of five.

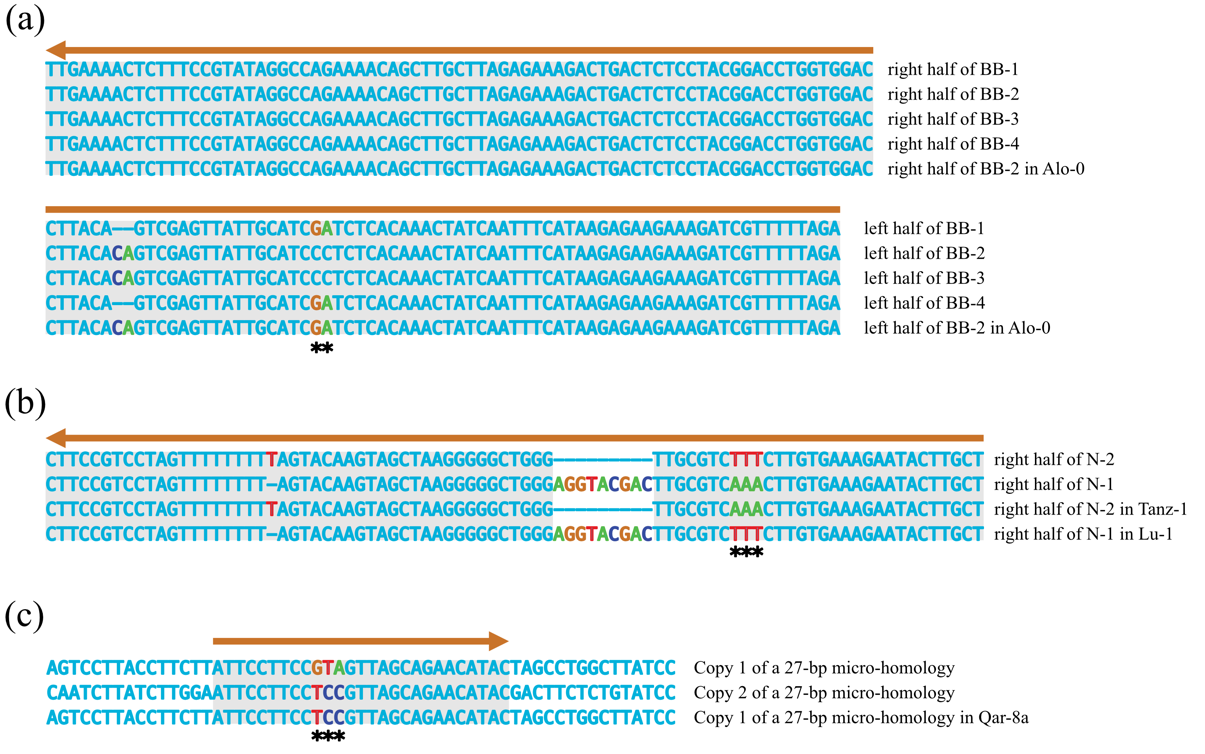

**Figure S10**. Examples of closely located small-scale variants explained by gene conversion events. (a-c) Sequences illustrate gene conversion events between different copies of non-tandem repeats. The asterisks indicate bases relating to potential gene conversion events.

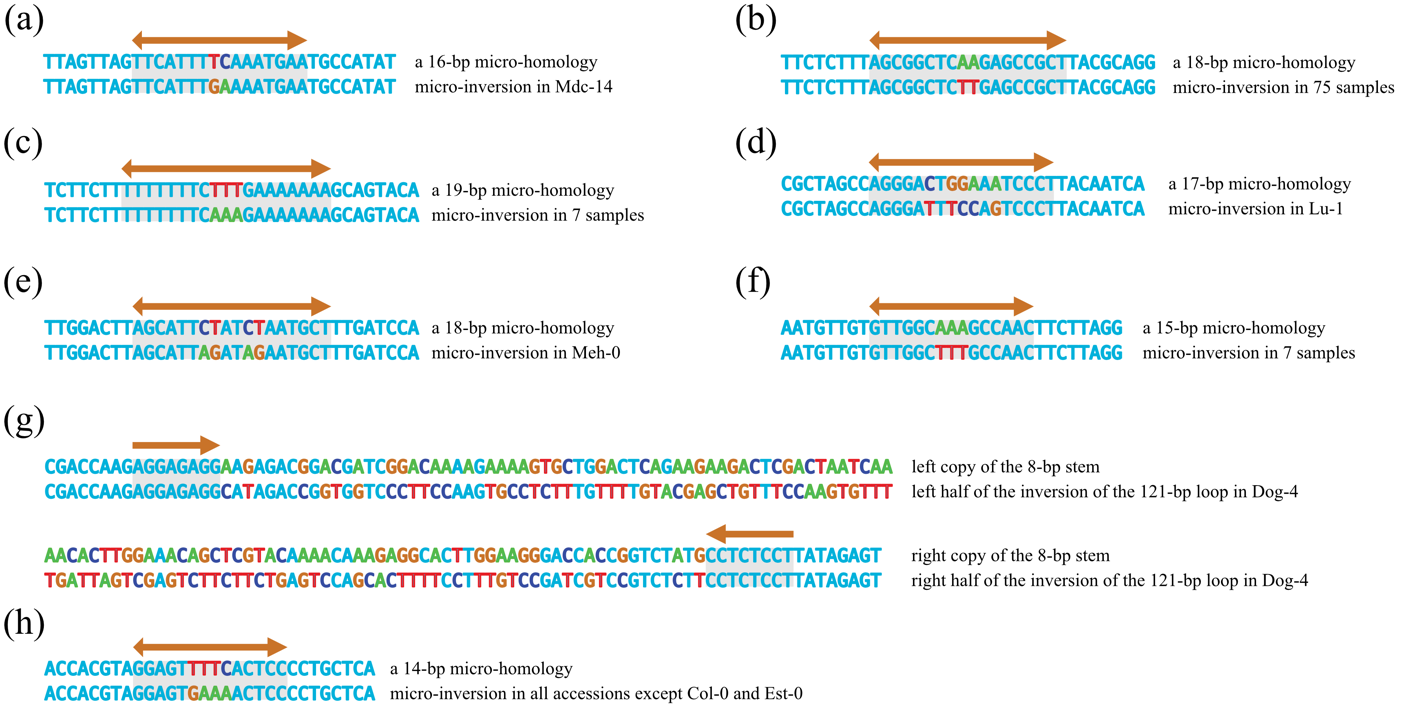

**Figure S11**. Examples of closely located small-scale variants explained by micro-inversion events in mt genomes. (a-h) Sequences illustrate by micro-inversion events mediated by partial palindromic repeats.

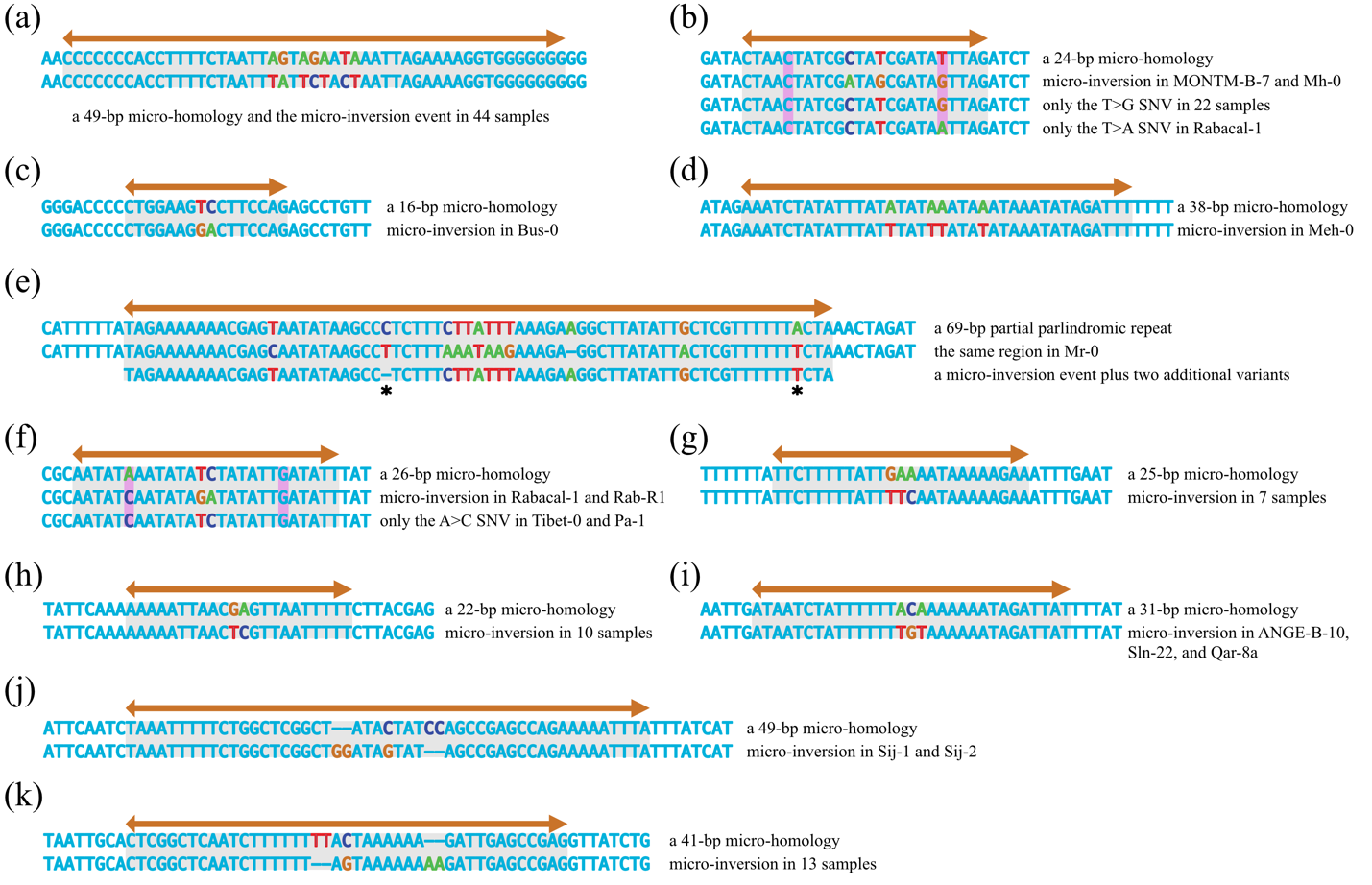

**Figure S12**. Examples of closely located small-scale variants explained by micro-inversion events in pt genomes. (a-k) Sequences illustrate by micro-inversion events mediated by partial palindromic repeats. The asterisks in panel e indicate bases relating to additional variants.

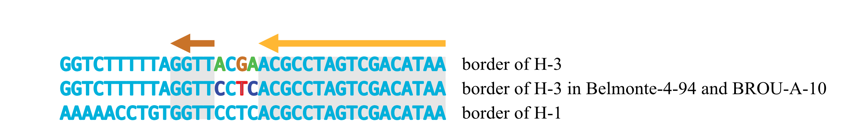

**Figure S13**. An example of closely located small-scale variants explained by repeat extension. Sequences illustrate a rare event involving the extension of the border repeat H-3 in two accessions.

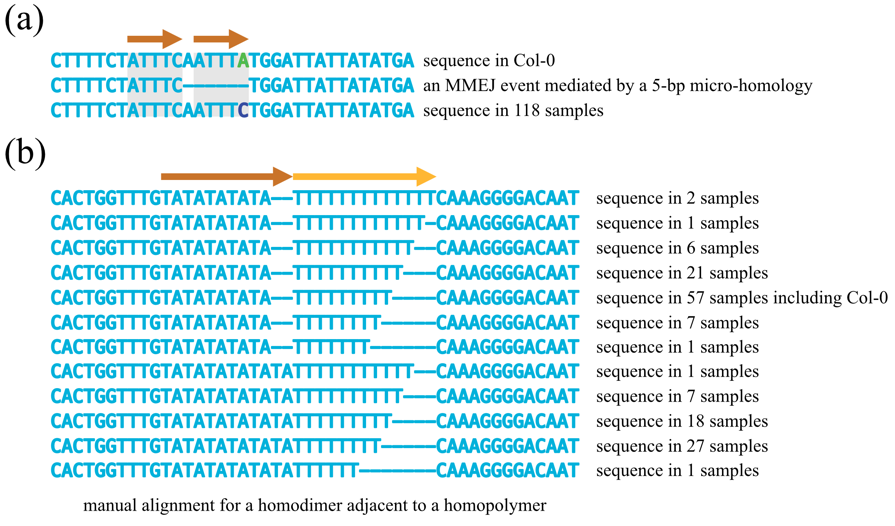

**Figure S14**. Some mutations were better explained when comparing multiple accessions. (a) An MMEJ event was interpreted as NHEJ because an additional SNV in Col-0. (b) Manual alignment was required when a homodimer was adjacent to a homopolymer.

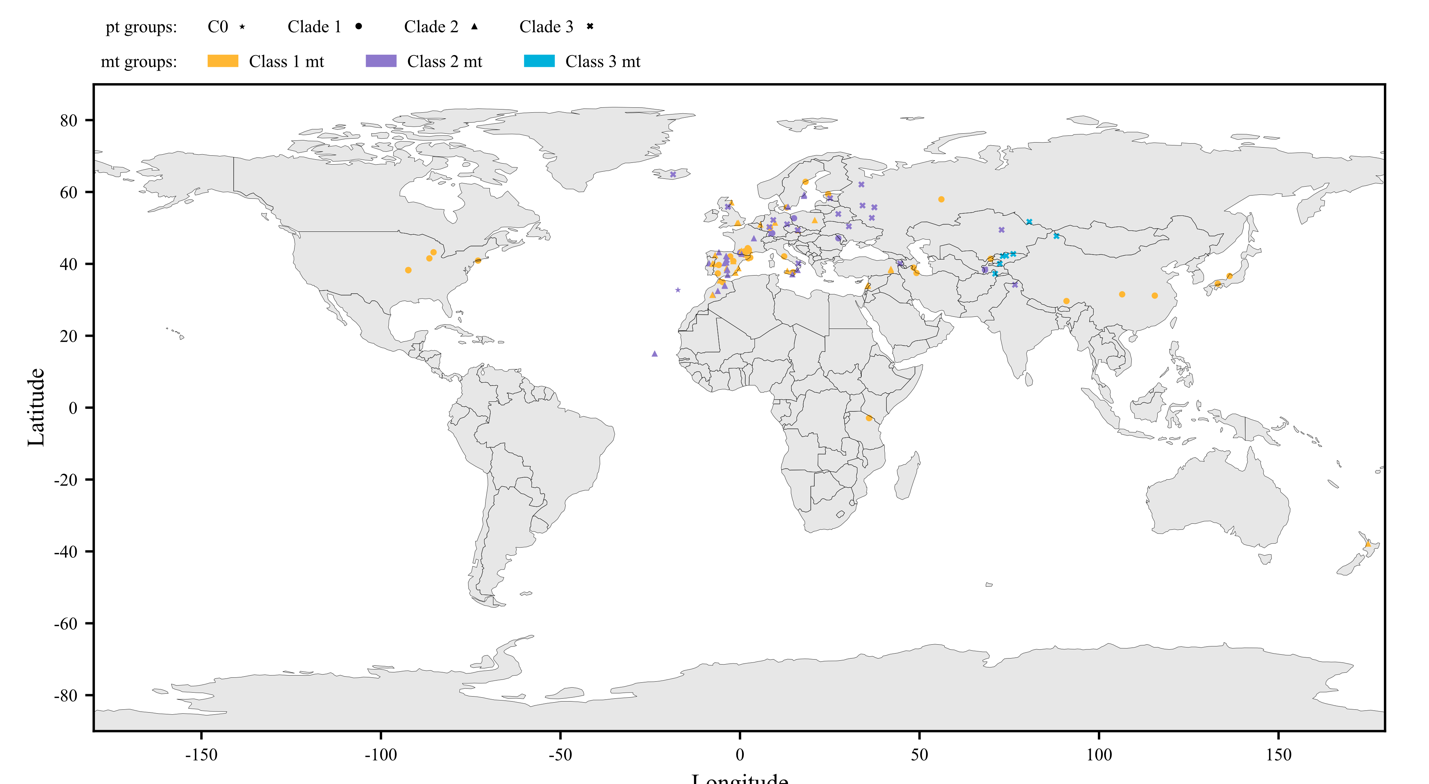

**Figure S15**. Geographic locations of the 149 samples used in this study. Clades in the pt SNV tree were indicated by shapes, while classes of mt genomes were indicated by colors.

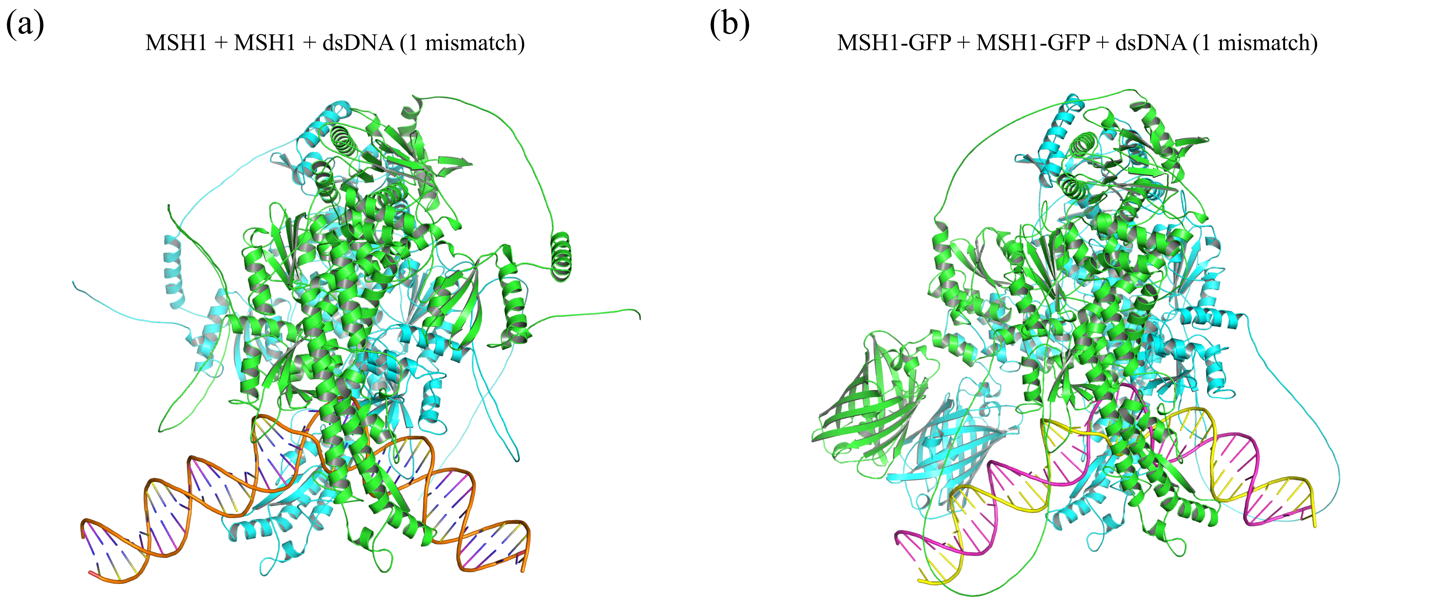

**Figure S16**. Structures of MSH1 dimers binding to double-stranded DNA with a mismatch. (a, b) AlphaFold 3 predicted that MSH1 dimers could bind to double-stranded DNA with a mismatch (b) when GFP was fused to its C-terminus, (a) or not.

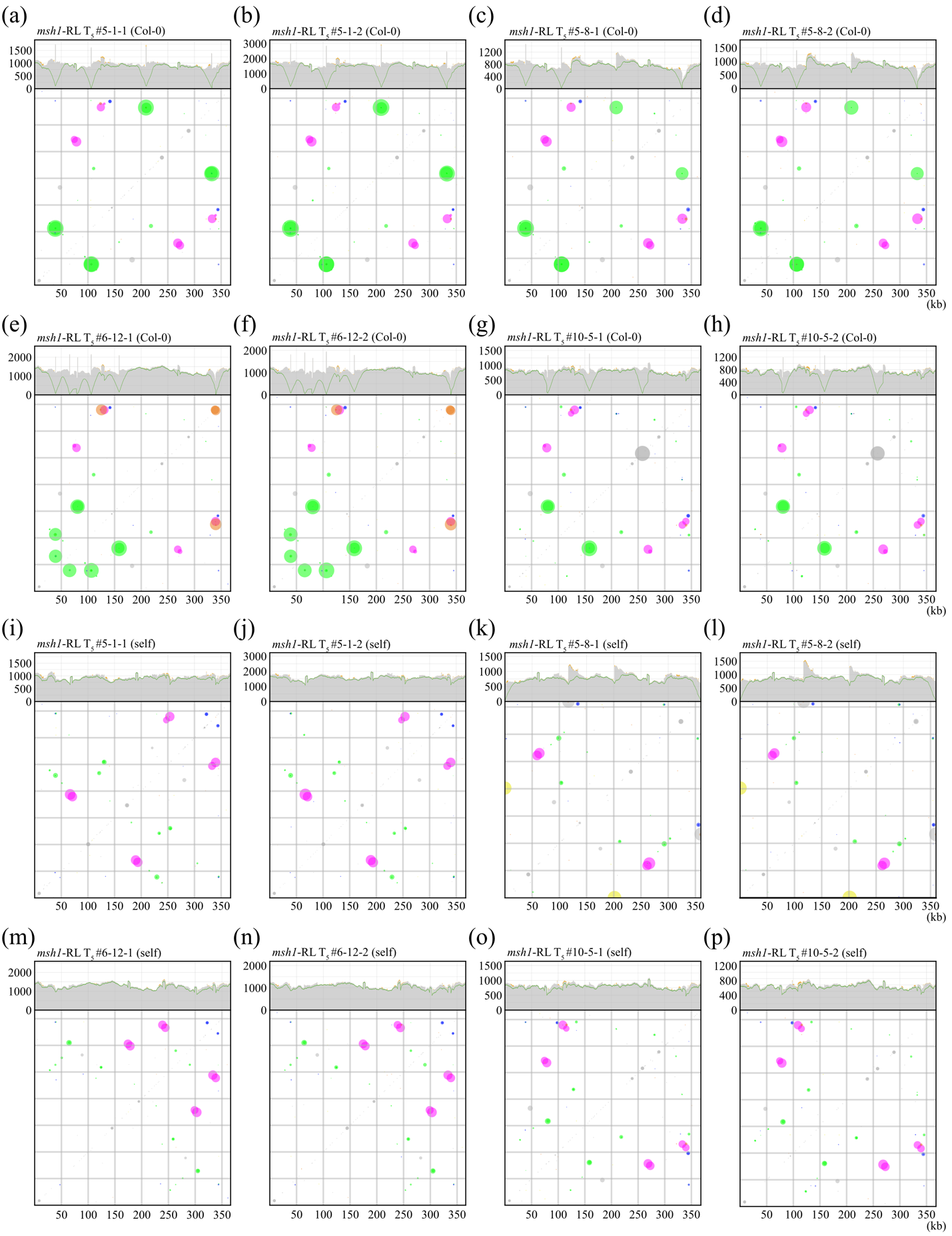

**Figure S17**. Coverages and SV frequencies in the mt genomes of *msh1* rescue lines. (a-p) Coverages (top) of HiFi reads from eight individuals of four *msh1* rescue lines mapped to mt genomes of (a-h) the original Col-0 reference, and (i-p) the newly assembled mt genome of the corresponding line. Grey areas indicate fully aligned reads, orange areas indicate partially aligned reads, and the green line indicate reads with no rearrangements plus one-rearrangement reads mediated by large repeats. Frequencies (bottom) of one-rearrangement reads in each sample. Colors represent categories of alignment overlap length (AO length) as in Zou et al. (2022).

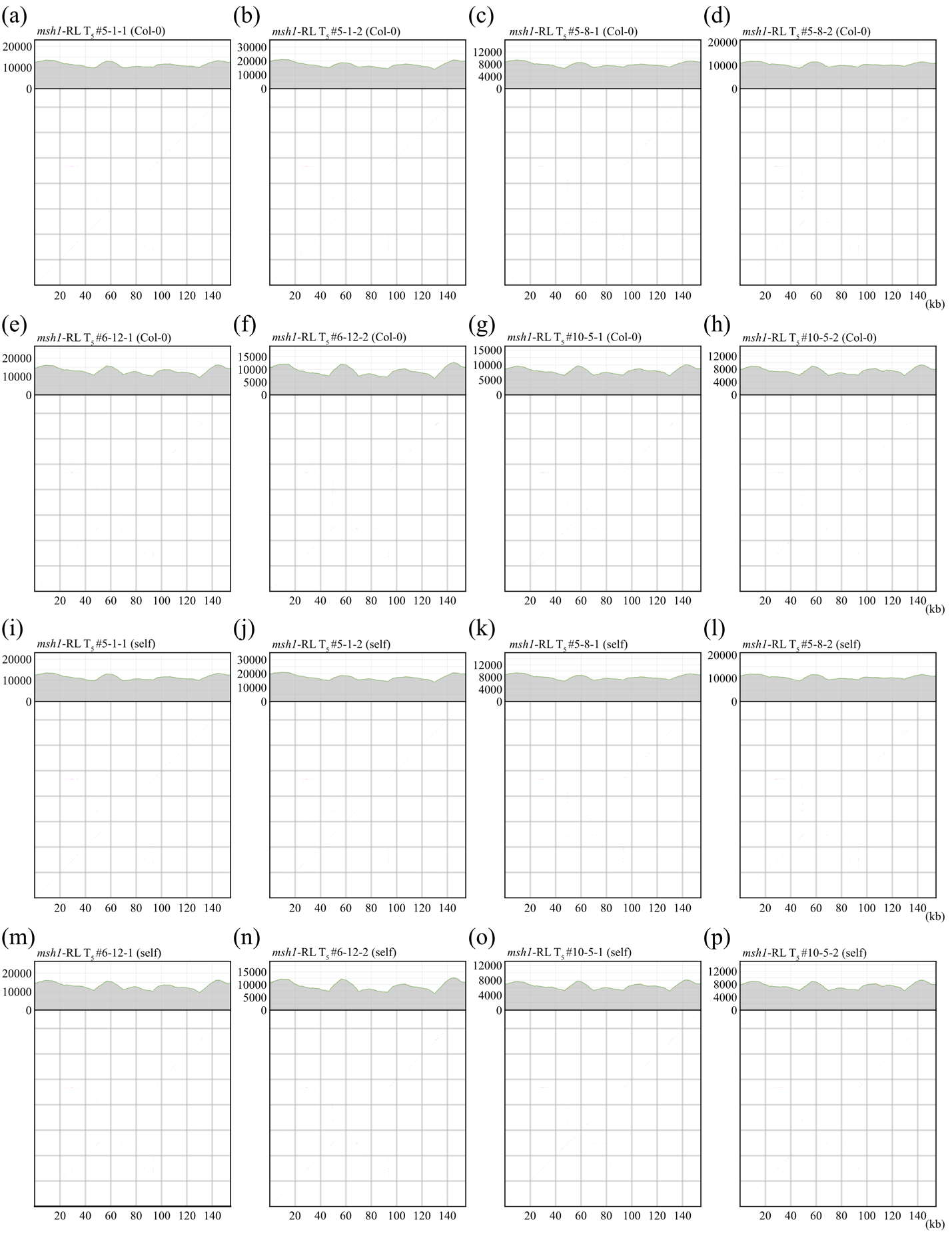

**Figure S18**. Coverages and SV frequencies in the pt genomes of *msh1* rescue lines. (a-p) Coverages (top) of HiFi reads from eight individuals of four *msh1* rescue lines mapped to pt genomes of (a-h) the original Col-0 reference, and (i-p) the newly assembled pt genome of the corresponding line. Grey areas indicate fully aligned reads, orange areas indicate partially aligned reads, and the green line indicate reads with no rearrangements plus one-rearrangement reads mediated by IRab. Frequencies (bottom) of one-rearrangement reads in each sample. Colors represent categories of alignment overlap length (AO length) as in Zou et al. (2022).

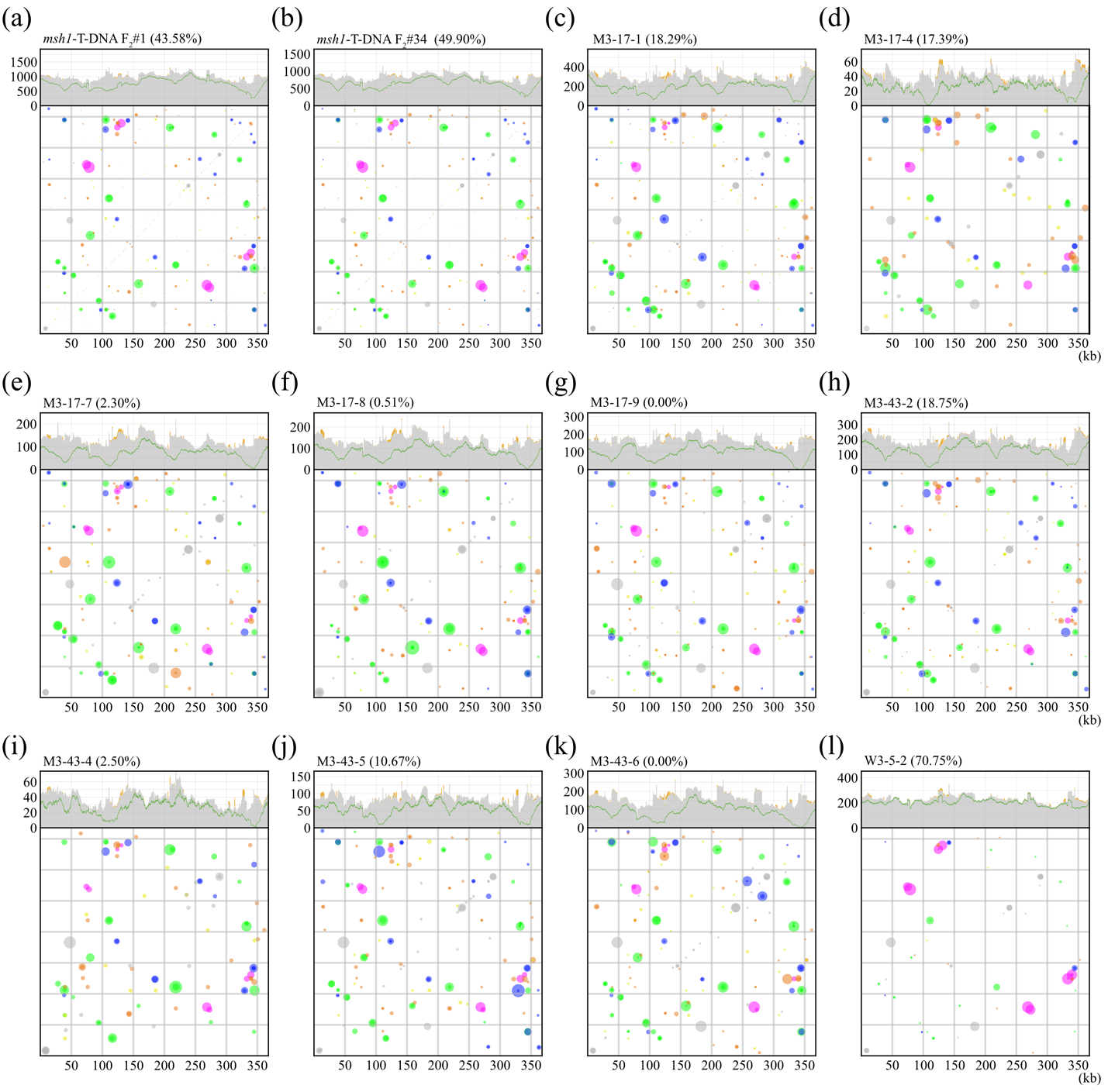

**Figure S19**. Coverages and SV frequencies in the mt genomes of *msh1* mutants. (a-l) Coverages (top) of HiFi reads from (a, b) two newly sequenced F_2_ individuals of *msh1* mutants, (c-k) eleven individuals (ten F_3_ *msh1* mutants and one wild type) from the published work (Zou et al., 2022). Grey areas indicate fully aligned reads, orange areas indicate partially aligned reads, and the green line indicate reads with no rearrangements plus one-rearrangement reads mediated by large repeats. Frequencies (bottom) of one-rearrangement reads in each sample. Colors represent categories of alignment overlap length (AO length) as in Zou et al. (2022). The percentage numbers in parentheses after the sample name indicate the lowest ratio of reference conformations around repeat Large1-2.

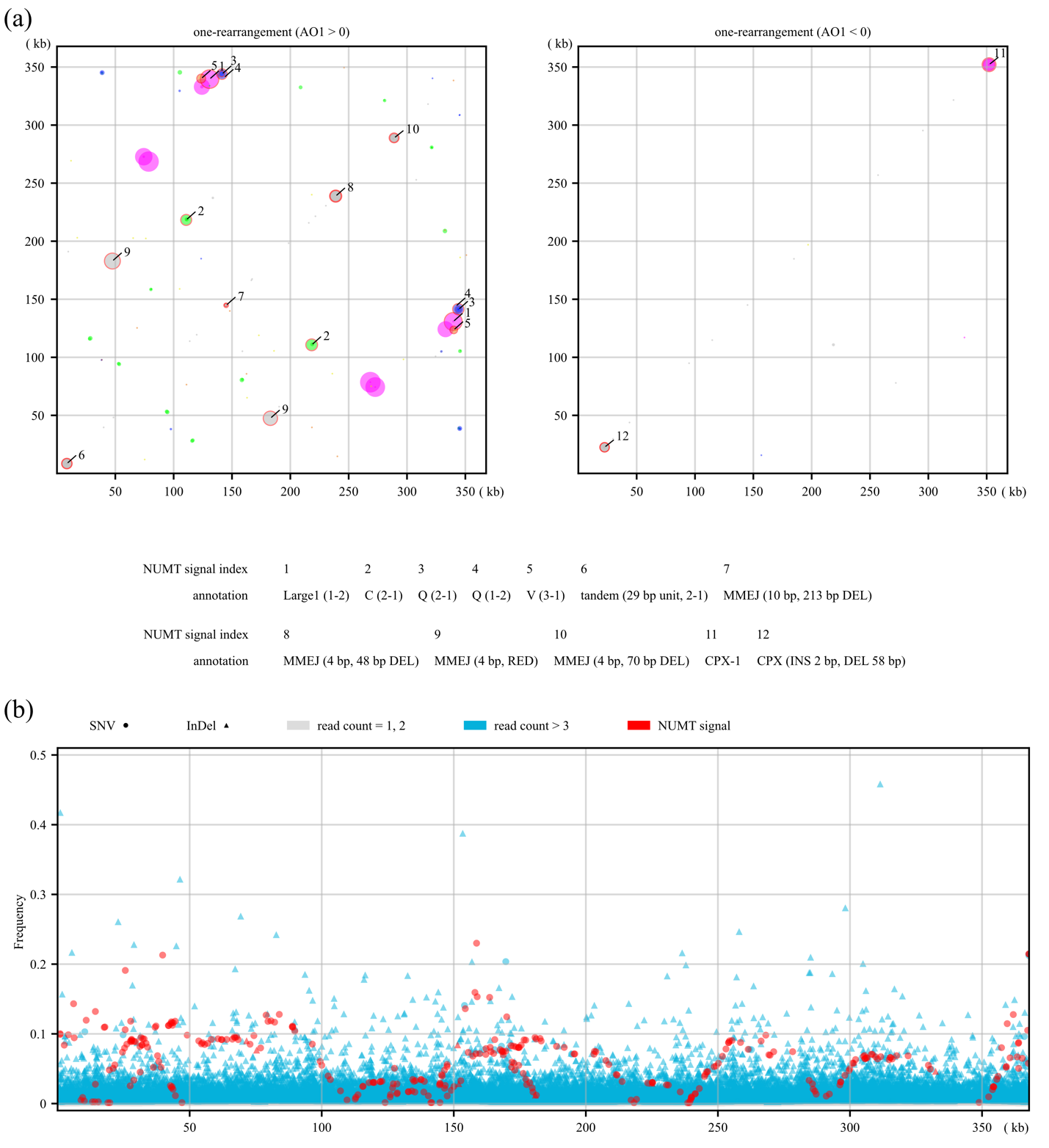

**Figure S20**. Frequencies of SVs and small-scale variants were affected by the giant NUMT on Chr2 of Col-0. (a) The frequences of twelve SVs were affected by the giant NUMT on Chr2 of Col-0. (b) The frequences of small-scale variants were affected by the giant NUMT on Chr2 of Col-0. Most of the affected variants were SNVs. The data from Col-CEN were used.

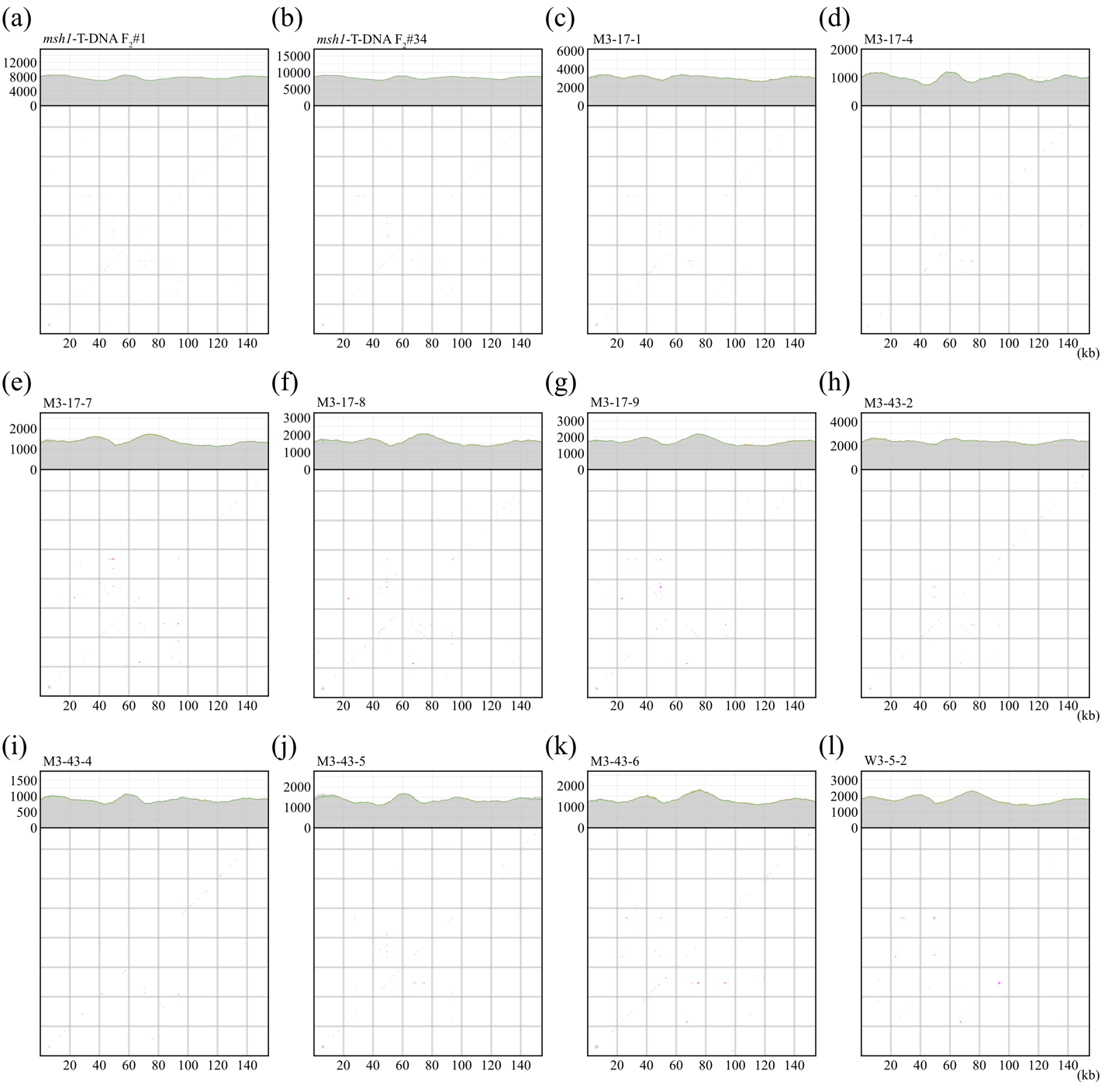

**Figure S21**. Coverages and SV frequencies in the pt genomes of *msh1* mutants. (a-l) Coverages (top) of HiFi reads from (a, b) two newly sequenced F_2_ individuals of *msh1* mutants, (c-k) eleven individuals (ten F_3_ *msh1* mutants and one wild type) from the published work (Zou et al., 2022). Grey areas indicate fully aligned reads, orange areas indicate partially aligned reads, and the green line indicate reads with no rearrangements plus one-rearrangement reads mediated by IRab. Frequencies (bottom) of one-rearrangement reads in each sample. Colors represent categories of alignment overlap length (AO length) as in Zou et al. (2022).

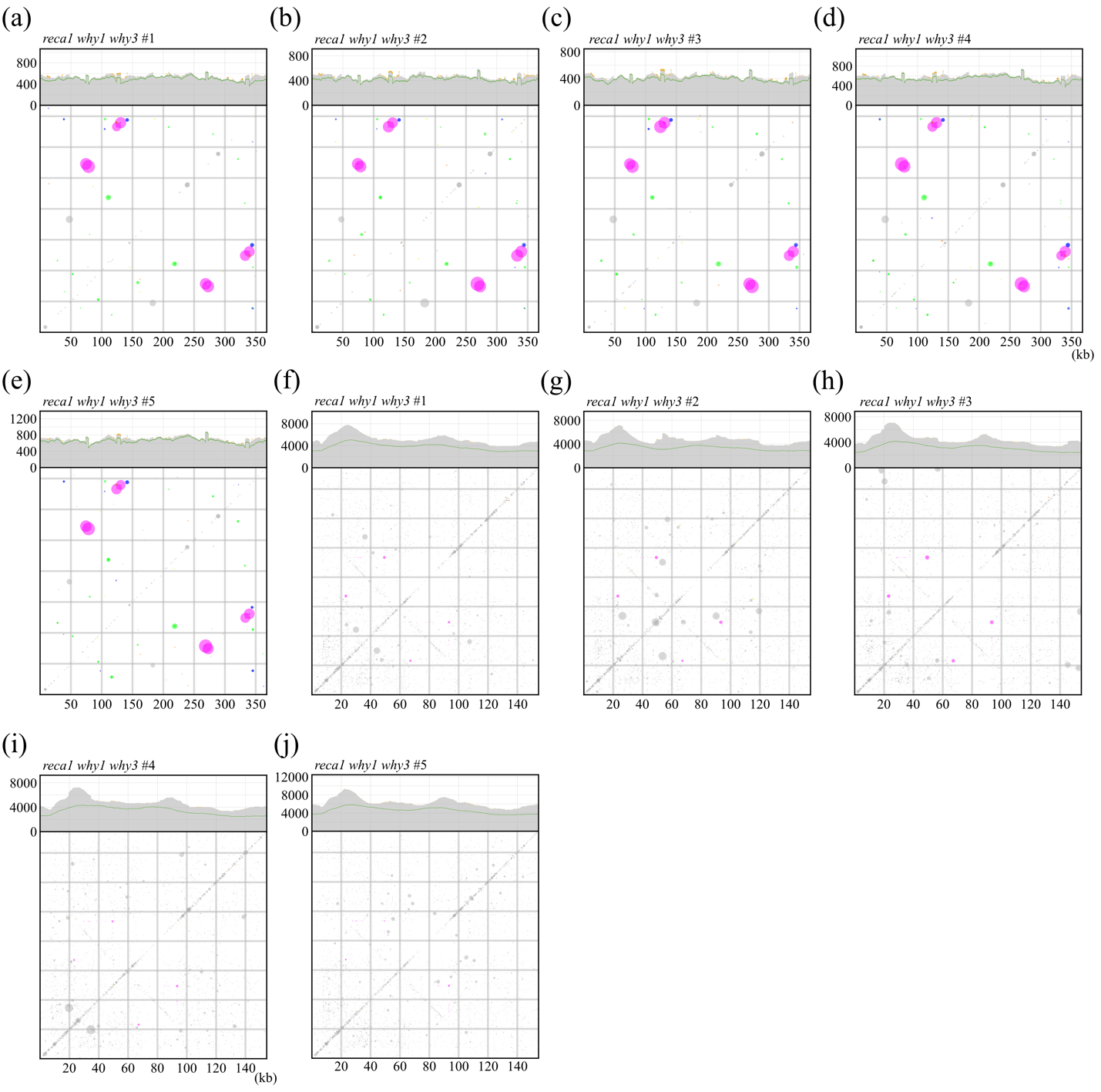

**Figure S22**. Coverages and SV frequencies in the mt and pt genomes of *recA1 why1 why3* triple mutants. (a-j) Coverages (top) of HiFi reads mapped to the Col-0 reference (a-e) mt and (f-j) pt genomes for five individuals of advanced-generation *recA1 why1 why3* triple mutants. Grey areas indicate fully aligned reads, orange areas indicate partially aligned reads, and the green line indicate reads with no rearrangements plus one-rearrangement reads mediated by large repeats (Large1, Large2, and IRab). Frequencies (bottom) of one-rearrangement reads in each sample. Colors represent categories of alignment overlap length (AO length) as in Zou et al. (2022).
